# Resolving artefacts in voltage-clamp experiments with computational modelling: an application to fast sodium current recordings

**DOI:** 10.1101/2024.07.23.604780

**Authors:** Chon Lok Lei, Alexander P. Clark, Michael Clerx, Siyu Wei, Meye Bloothooft, Teun P. de Boer, David J. Christini, Trine Krogh-Madsen, Gary R. Mirams

## Abstract

Cellular electrophysiology is the foundation of many fields, from basic science in neurology, cardiology, oncology to safety critical applications for drug safety testing, clinical phenotyping, etc. Patch-clamp voltage clamp is the gold standard technique for studying cellular electrophysiology. Yet, the quality of these experiments is not always transparent, which may lead to erroneous conclusions for studies and applications. Here, we have developed a new computational approach that allows us to explain and predict the experimental artefacts in voltage-clamp experiments. The computational model captures the experimental procedure and its inadequacies, including: voltage offset, series resistance, membrane capacitance and (imperfect) amplifier compensations, such as series resistance compensation and supercharging. The computational model was validated through a series of electrical model cell experiments. Using this computational approach, the artefacts in voltage-clamp experiments of cardiac fast sodium current, one of the most challenging currents to voltage clamp, were able to be resolved and explained through coupling the observed current and the simulated membrane voltage, including some typically observed shifts and delays in the recorded currents. We further demonstrated that the typical way of averaging data for current-voltage relationships would lead to biases in the peak current and shifts in the peak voltage, and such biases can be in the same order of magnitude as those differences reported for disease-causing mutations. Therefore, the presented new computational pipeline will provide a new standard of assessing the voltage-clamp experiments and interpreting the experimental data, which may be able to rectify and provide a better understanding of ion channel mutations and other related applications.

## 1 Introduction

The pioneering work of Hodgkin and Huxley (1939, 1952) provided fundamental insights into cellular excitability, and marked the start of the modern era of systems electrophysiology research — this field has provided deep understanding of diverse biological processes in the areas of neurology, cardiology, oncology, safety critical applications for drug safety testing, and much more. Patch-clamp experiments have been the gold standard for the study of cellular electrophysiology since their invention (Sakmann and Neher, 1984). In particular, whole-cell mode voltage clamp allows us to probe the dynamic current responses of an electrically excitable cell while controlling its voltage. Measured whole-cell current is typically carried by a collection of numerous transmembrane ion-permeable proteins, each with distinct opening and closing kinetics (often nonlinear) that are studied through pharmacological block and step-wise voltage experiments. These studies and applications rely on the assumption that membrane voltage is *perfectly* clamped at desired voltages. Here, we aim to demonstrate when and how this assumption breaks down, and provide strategies to improve interpretation of voltage-clamp data.

Imperfect voltage clamping is a pervasive issue in electrophysiology research and contributes to experimental artefacts that cause incorrect characterisation of cellular properties. Errors primarily arise due to cell and pipette capacitances, leak currents, and the series resistance between the pipette tip and cell membrane (Marty and Neher, 1983).

Efforts have been devoted to develop strategies to correct for artefact effects over the last few decades. In post-processing, Marty and Neher (1983) suggested a correction to current-voltage (I-V) summary curves: using estimates of the series resistance to predict voltage drop for a given current, and then to translate each point horizontally on the I-V curve appropriately. But this approach does not account for all the artefacts we know to be present in the system, or correct for the loss of voltage clamp that can occur during an experiment. Modern amplifiers have compensatory mechanisms designed to *partially* correct for the unwanted effects of artefacts during acquisition. Cell capacitance is compensated using a mechanism that speeds membrane charging (Sigworth, 1995a; Sigworth et al., 1995). Amplifiers also compensate for artefacts that result from series resistance: namely, the slowing of the membrane voltage’s approach to the command voltage (Sigworth et al., 1995; Armstrong and Chow, 1987), and the deviation of the membrane voltage from the command voltage (Moore et al., 1984; Strickholm, 1995; Sherman et al., 1999; Weerakoon et al., 2009).

These strategies, however, still rely on imperfect estimates of series resistance and capacitance, and create a system that is sensitive to overcompensation, with unphysiological dynamics. Thus, whole cell patch-clamp artefacts remain an issue, especially when measuring currents with fast dynamics, such as sodium.

While it has long been known that series resistance skews current-voltage curves (Marty and Neher, 1983), computational modelling of artefacts to assist in data interpretation is a relatively new approach (Lei et al., 2020a; Lei, 2020; Montnach et al., 2021; Abrasheva et al., 2024). We showed that a series-resistance compensation model could improve fits to potassium channel data (Lei et al., 2020a). Montnach et al. (2021) showed how a similar approach improved interpretation of recorded sodium dynamics. Lei (2020) developed a computational model of the amplifier dynamics compensating for the slow approach of membrane voltage to the command voltage — a compensation technique called *prediction* or *super-charging*, and Abrasheva et al. (2024) showed how the supercharging aspect improves ion current model fits to sodium data however potentially failing to correct the steady state due to missing series resistance compensation in their model.

In this study, we develop a new computational approach that allows us to explain and predict artefacts during voltage-clamp experiments and further improve interpretation of sodium current recordings (Fig. 1). The computational model includes equations that comprehensively reproduce voltage-clamp artefacts and the compensatory circuits in amplifiers (e.g., capacitances/series resistance compensation, supercharging), including separate parameters for estimated and true values of series resistance and cell capacitance. We validate and test the model on sodium current at physiological temperature, with multiple patching strategies/cell types, and several compensation levels to demonstrate the artefact model’s ability to reproduce dynamics under extreme and varying conditions. We show the ability of this computational model to reproduce artefact-causing dynamics in an electronic model cell, in human induced pluripotent stem cell-derived cardiomyocytes (hiPSC-CMs), and in Na_V_1.5-expressing human embryonic kidney (HEK) cells. The model improves the interpretation of data recorded with different amplifiers, using rupture or perforated patch, and both manual and automated patch techniques. We believe this model represents a step forward in how voltage-clamp data can be interpreted. Specifically, we (1) provide a tool to improve interpretation of electrophysiological recordings, and (2) make it possible to obtain accurate estimates of biophysical properties that were previously obscured by experimental artefacts.

**Figure 1:**
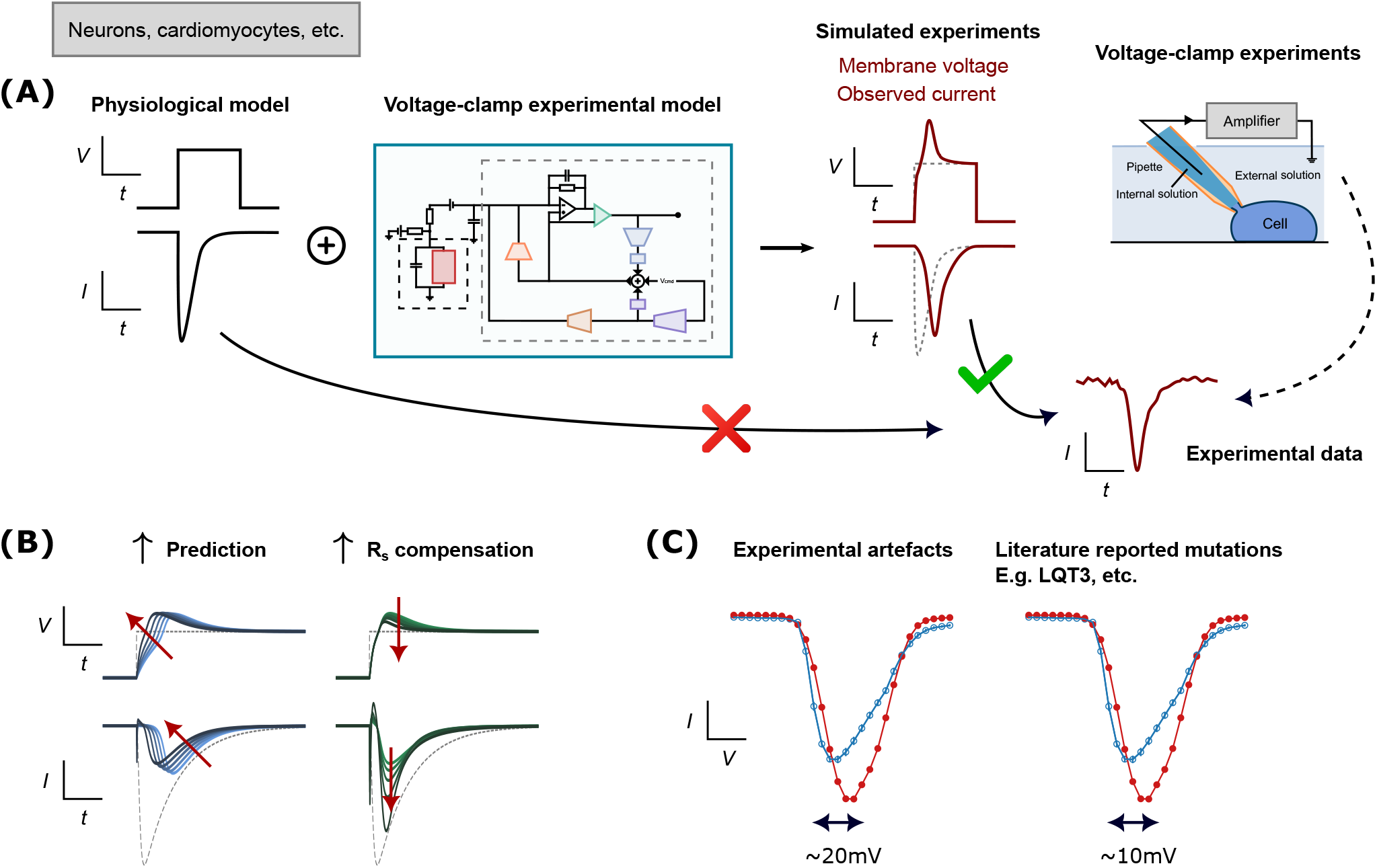
A schematic overview of the study. **(A)** A workflow schematic that shows by combining a physiological model with our mathematical model of patch-clamp voltage clamp (Fig. 2), it produces simulated experimental data that can be *correctly* compared to the experimental data. **(B)** Our new model explains the effects of different types of compensation. **(C)** Experimental artefacts may result in error that is similar to or even larger than differences due to mutations reported in the literature.

## 2 Results

### 2.1 A detailed computational model for patch-clamp voltage-clamp experiments

The first version of the mathematical model of voltage-clamp experimental artefacts was published in Lei et al. (2020a); here we expand it to include all the essential functionalities of a modern voltage-clamp amplifier. The new supercharging component of our model was introduced in the first-author’s PhD thesis (Lei, 2020) and independently reported by Abrasheva et al. (2024). Fig. 2 shows a schematic of the equivalent electrical circuit of the new detailed model for the whole-cell patch-clamp voltage clamp experiment, including the major commonly-used compensation methods. The new model equations are:

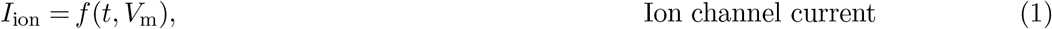

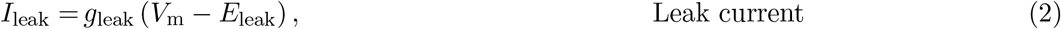

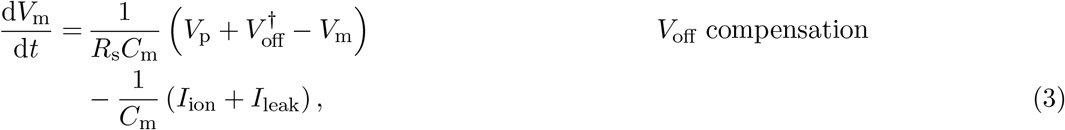

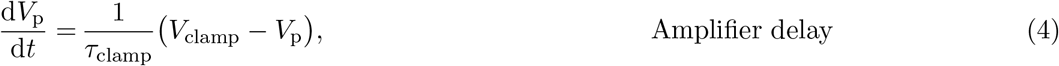

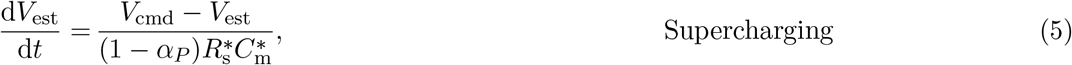

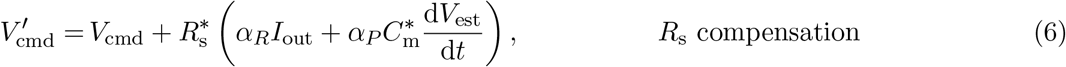

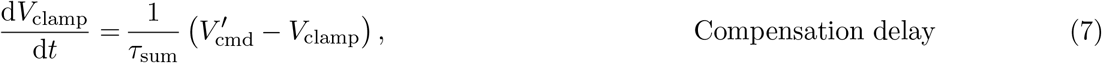

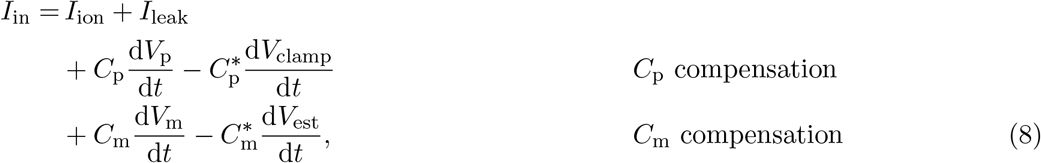

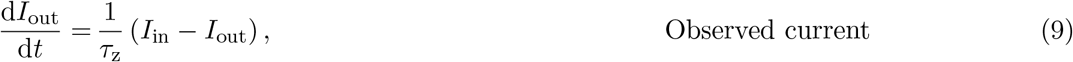

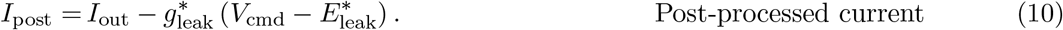

**Figure 2:**
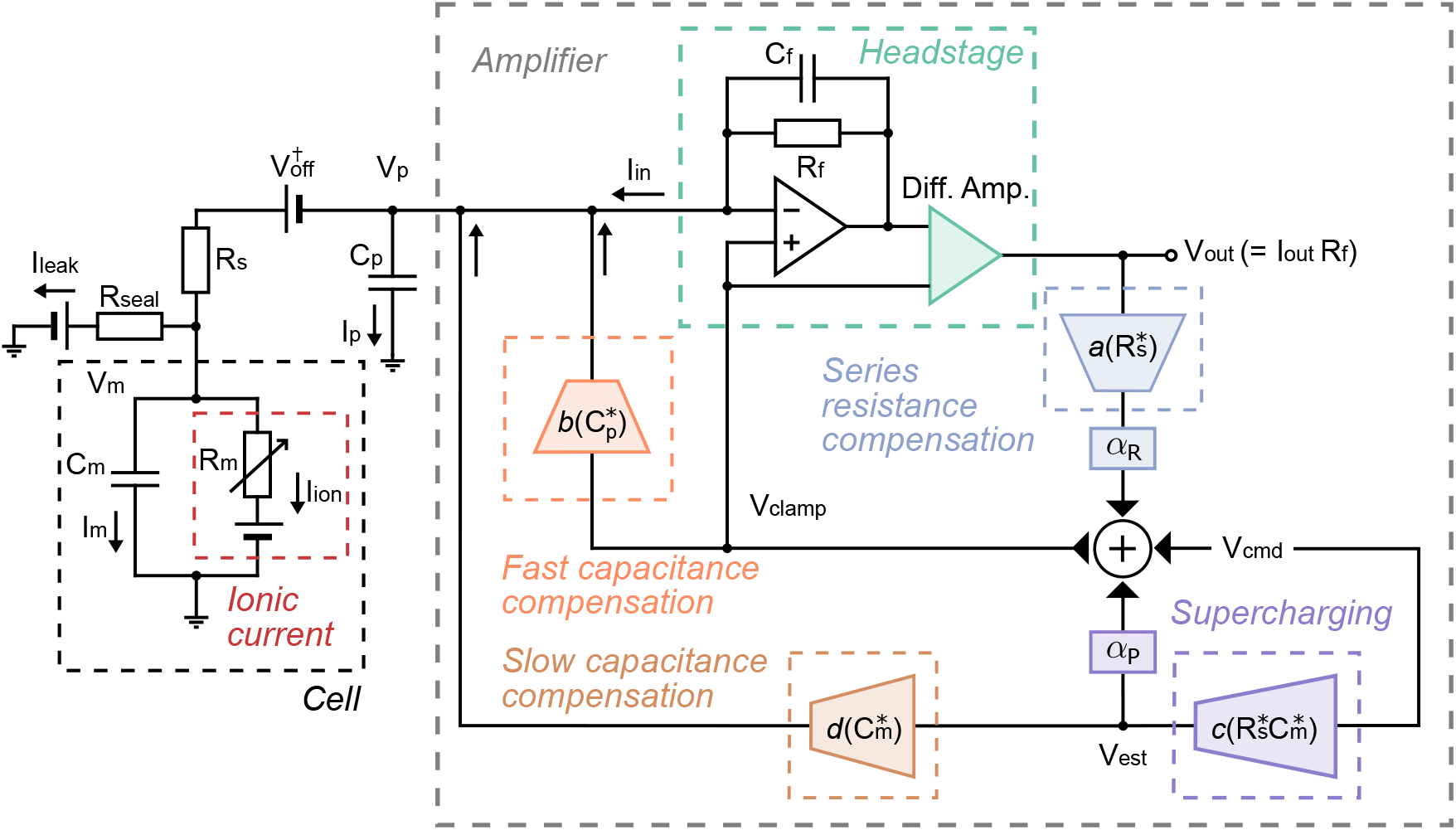
A schematic of the equivalent electrical circuit of the new detailed mathematical model for whole-cell patch-clamp voltage-clamp experiments. Right shows the components of an patch-clamp amplifier, connecting to the cell on the left. The amplifier includes the components for compensating for series resistance, supercharging, slow capacitance, fast capacitance, and voltage offset. The leak correction is modelled as a post-processing step that is not part of the equivalent circuit.

An explanation of the symbols and parameters is provided in Table 1. The model quantitatively describes the dynamic interaction between the voltages and currents in the experimental system, as well as the effects of amplifier compensations: *α*_*R*_ is series resistance compensation level, and *α*_*P*_ is the supercharging compensation (or prediction) level. We will refer to this set of equations as the *voltage-clamp model*.

**Table 1:**
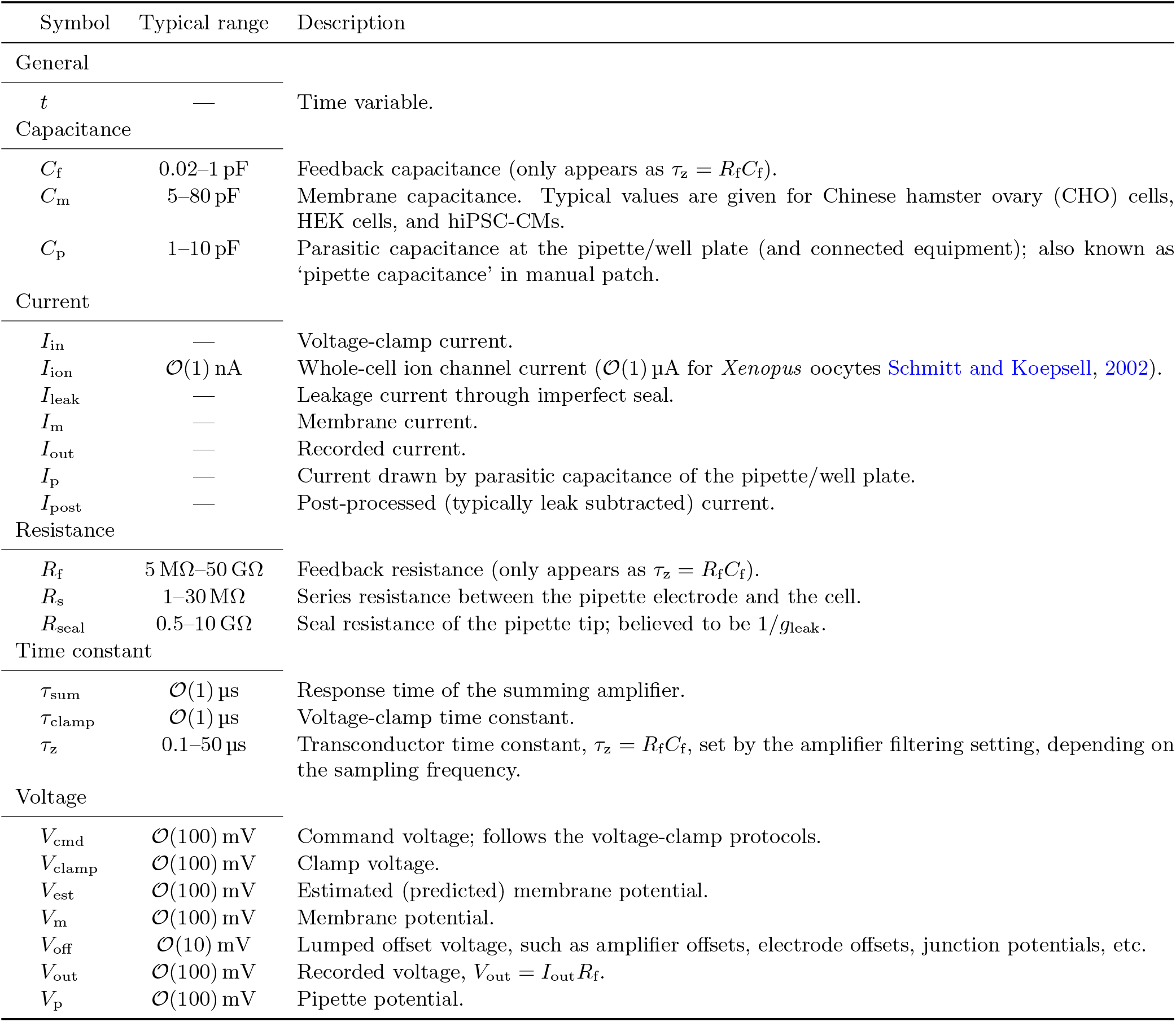
Glossary of symbols and parameters. A machine or post-processing estimate of a parameter *X* is denoted as *X*^*\**^, and the error in the estimate of the same parameter as *X*^*†*^. The values are taken from Neher (1995); HEKA Elektronik GmbH (2018); Axon Instruments Inc. (1999), unless otherwise specified.

| Symbol | Typical range | Description |
| --- | --- | --- |
| General |  |  |
| $t$ | — | Time variable. |
| Capacitance |  |  |
| $C_f$ | 0.02–1 pF | Feedback capacitance (only appears as $\tau_z = R_f C_f$). |
| $C_m$ | 5–80 pF | Membrane capacitance. Typical values are given for Chinese hamster ovary (CHO) cells, HEK cells, and hiPSC-CMs. |
| $C_p$ | 1–10 pF | Parasitic capacitance at the pipette/well plate (and connected equipment); also known as ‘pipette capacitance’ in manual patch. |
| Current |  |  |
| $I_{in}$ | — | Voltage-clamp current. |
| $I_{ion}$ | $\mathcal{O}(1)$ nA | Whole-cell ion channel current ($\mathcal{O}(1)$ μA for *Xenopus* oocytes Schmitt and Koepsell, 2002). |
| $I_{leak}$ | — | Leakage current through imperfect seal. |
| $I_m$ | — | Membrane current. |
| $I_{out}$ | — | Recorded current. |
| $I_p$ | — | Current drawn by parasitic capacitance of the pipette/well plate. |
| $I_{post}$ | — | Post-processed (typically leak subtracted) current. |
| Resistance |  |  |
| $R_f$ | 5 MΩ–50 GΩ | Feedback resistance (only appears as $\tau_z = R_f C_f$). |
| $R_s$ | 1–30 MΩ | Series resistance between the pipette electrode and the cell. |
| $R_{seal}$ | 0.5–10 GΩ | Seal resistance of the pipette tip; believed to be $1/g_{leak}$. |
| Time constant |  |  |
| $\tau_{sum}$ | $\mathcal{O}(1)$ μs | Response time of the summing amplifier. |
| $\tau_{clamp}$ | $\mathcal{O}(1)$ μs | Voltage-clamp time constant. |
| $\tau_z$ | 0.1–50 μs | Transconductor time constant, $\tau_z = R_f C_f$, set by the amplifier filtering setting, depending on the sampling frequency. |
| Voltage |  |  |
| $V_{cmd}$ | $\mathcal{O}(100)$ mV | Command voltage; follows the voltage-clamp protocols. |
| $V_{clamp}$ | $\mathcal{O}(100)$ mV | Clamp voltage. |
| $V_{est}$ | $\mathcal{O}(100)$ mV | Estimated (predicted) membrane potential. |
| $V_m$ | $\mathcal{O}(100)$ mV | Membrane potential. |
| $V_{off}$ | $\mathcal{O}(10)$ mV | Lumped offset voltage, such as amplifier offsets, electrode offsets, junction potentials, etc. |
| $V_{out}$ | $\mathcal{O}(100)$ mV | Recorded voltage, $V_{out} = I_{out} R_f$. |
| $V_p$ | $\mathcal{O}(100)$ mV | Pipette potential. |

### 2.2 Experimental validation of the patch-clamp voltage-clamp model

The new mathematical model of patch-clamp voltage clamp was validated using a tailor-made electrical circuit — a “model cell” — connected to the voltage-clamp amplifier as one normally would connect a biological cell (see Supplementary Materials for the full setup). The model cell was designed with multiple resistor-capacitor (RC) circuits in series and in parallel to elicit current responses similar to a biological cell. It was then connected with an explicit series resistor and capacitor before connecting to the amplifier, emulating the real experimental conditions in patch clamping. In this set-up, the model cell’s “membrane voltage” can be measured directly using a second amplifier running in current-clamp mode (with zero current injection).

Below, we derive the equations of the “ionic current” (*I*_ion_) for the electrical model cell. By analysing the electrical components with Kirchhoff’s circuit laws, we obtain

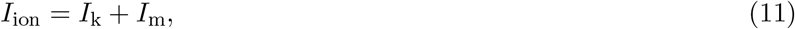

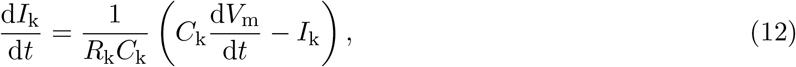

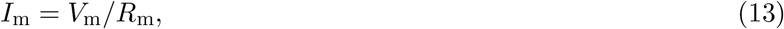

where the values of the model cell components are given in Fig. 3. The choice of the time constant of the dynamical part of the current, given by the product *R*_k_*C*_k_ = 1 ms, ensures the correct time scale for testing our new model, including the effects of the supercharging (prediction) component; we note that the dynamics of the overall current also depends on the dynamics of membrane voltage (*V*_m_). To mimic the effect of large currents such as the fast sodium current, the overall current magnitude of *I*_ion_ of the model cell is *𝒪* (10) nA. With these choices, the model cell experiments provide a strong ‘stress-test’ for the mathematical model of voltage clamp.

**Figure 3:**
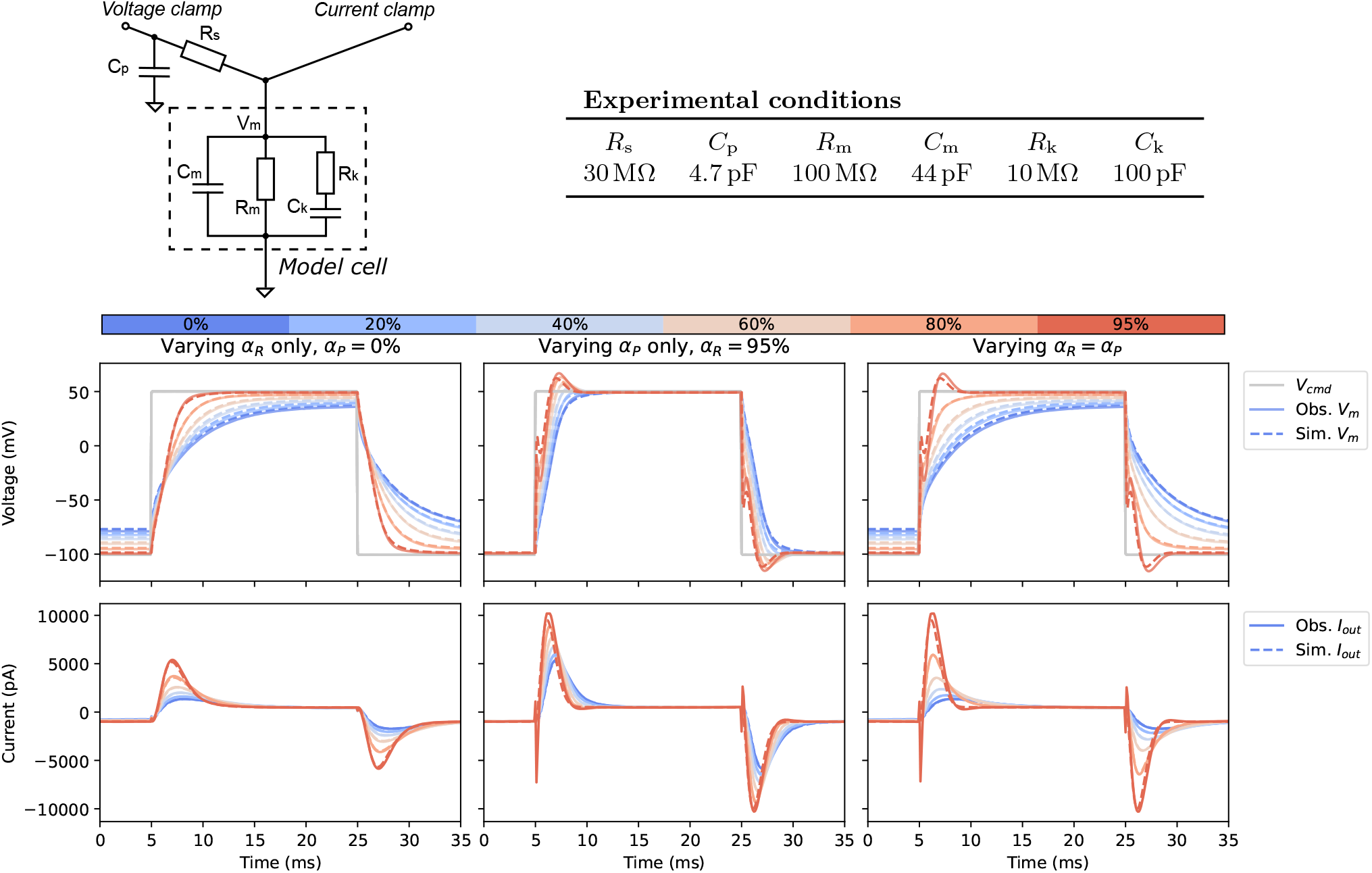
Electrical model-cell validation experiments. **Top:** An electrical schematic of the model cell (a modification of the ‘Type II’ model cell used in Lei et al. (2020a) with faster dynamics) and the chosen nominal component values. **Bottom: (Left)** Effect of series-resistance compensation (*α*_*R*_); **(middle)** supercharging compensation (*α*_*P*_); and **(right)** varying both compensation levels together. Grey solid/dashed lines are the measured/simulated command voltage *V*_cmd_. Coloured solid/dashed lines are the measured/simulated **(top)** membrane voltage *V*_m_ and **(bottom)** output current *I*_out_ at different levels of series resistance compensation (*α*_*R*_) and supercharging (*α*_*P*_).

Fig. 3 shows the validation results of the voltage-clamp model via our electrical model cell experiments. Typically, the highest recommended compensation is around 80 %, but to perform stress tests of the system and the model, we pushed the amplifier compensation to a maximum of 95 %; whether the maximum possible value can be set to 95 % was actually dependent on the dynamics of the current of interest and how it coupled with the built-in filtering and feedback of the amplifier. We note that such a high level of compensation is generally *not* recommended for biological cell experiments, as the oscillations of potential may destroy the cells.

Three types of compensation settings were tested: For the first set (left of Fig. 3), the supercharging (prediction, *α*_*P*_) was fixed to zero and series resistance compensation (*α*_*R*_) was varied from 0 % (no compensation) to 80 % with an increment of 20 %, and 95 %. Setting *α*_*P*_ = 0 reduces the model to that presented in Lei et al. (2020a) i.e. without supercharging. In the second set (middle), we tested when *α*_*R*_ was set to the maximum (95 %) and *α*_*P*_ varied from 0 % to its maximum. Finally (right), we tested when *α*_*R*_ and *α*_*P*_ were varied together. This last condition is most commonly used during biological experiments.

A simple large voltage step was used as the command voltage, as it should elicit relatively large current from the model cell, mimicking the current and artefact sizes seen in fast sodium current recordings. The mathematical model (dashed lines in Fig. 3) was able to predict the experimental results (solid lines) excellently for compensation levels up to 80 %, with differences generally not visible to the eye in Fig. 3. For 95 % compensation of both series resistance and prediction, although the simulations become distinguishable from the measurements, the qualitative shapes of the responses are correct with complex deflections, overshoots and turning points all predicted well. Therefore, most importantly, these validation results showed that as long as we have the correct model for the ionic current, we are able to predict and correct for experimental errors well.

### 2.3 Computational model predicts compensation effects on *in vitro* fast sodium current measurements

We next explored whether the computational voltage-clamp model could predict the effect of changes in compensation levels on experimental recordings of the fast sodium current *in vitro*. Since easily distinguishable effects when varying compensation levels were desired, the perforated patch technique was employed to induce relatively high series/access resistance. Experiments were performed in hiPSC-CMs at 37 °C; further details are given in **Methods**.

The fast sodium current *in vitro* model was used here to elucidate how artefacts can contaminate and distort the underlying physiological current in the recordings. Fig. 4 shows *in vitro* assessments of our voltage-clamp experimental artefact model when applied to a fast sodium current model (Paci et al., 2020, left), and as a reference, the ‘ideal’ voltage clamp where no experimental artefacts are included is shown in black dashed lines (i.e. when Eq. (1) was simulated directly with *V* cmd as input). The results were compared to *in vitro* patch-clamp voltage-clamp experimental data measured from hiPSC-CMs (Fig. 4 right).

**Figure 4:**
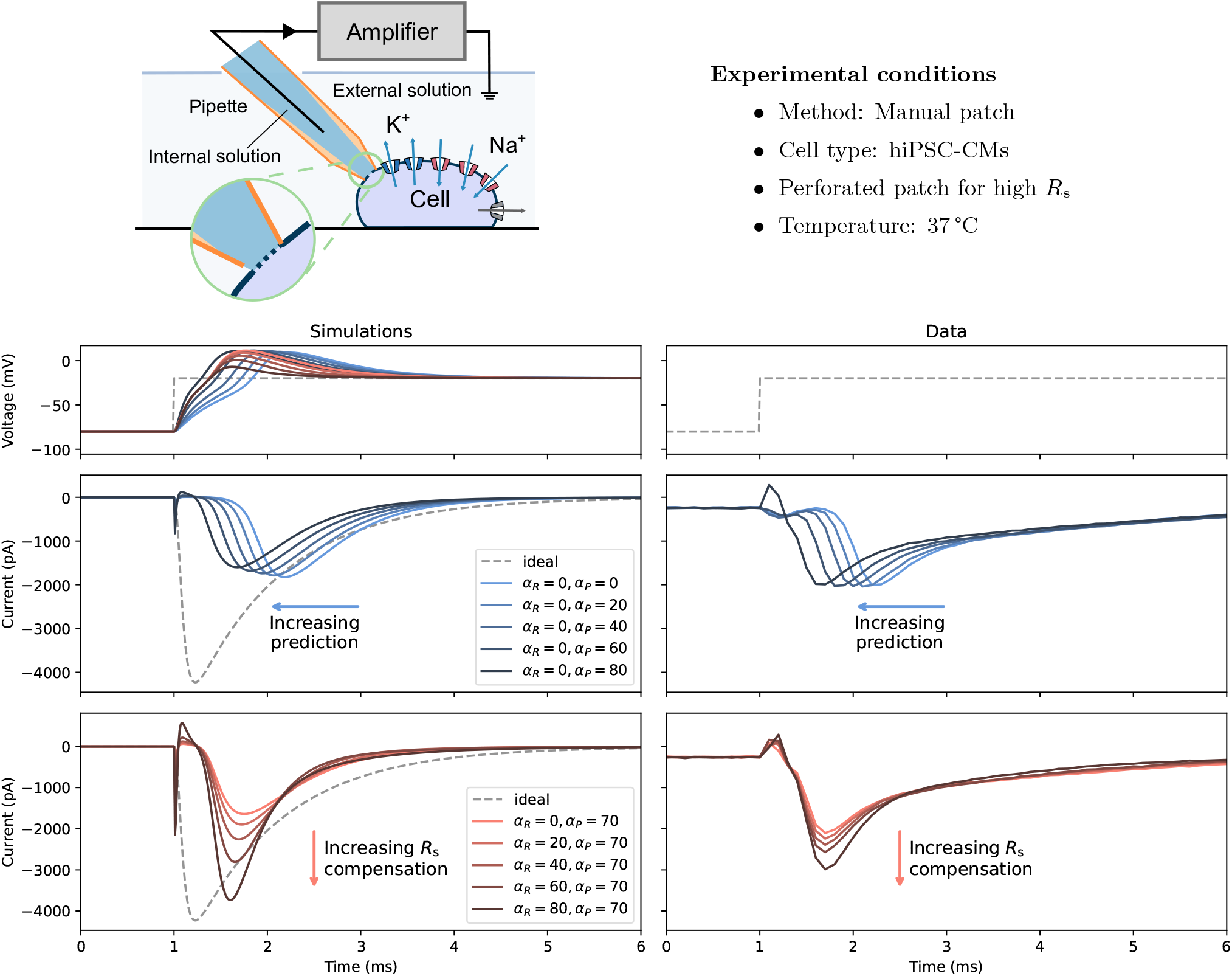
Assessing fast sodium current (*I*_Na_) responses to experimental artefacts, with recordings in hiPSC-CMs as an example. **Top:** A schematic of the experimental setup and the experimental conditions for the perforated patch experiments with hiPSC-CMs. **Bottom: (Left)** Simulations of the voltage-clamp model (Equations (1)–(10)) using a computational hiPSC-CM model for I_ion_; **(Right)** Patch-clamp voltage-clamp *in vitro* experimental data measured from hiPSC-CMs. From top to bottom, it shows the command voltage (dashed lines) and the membrane voltage (solid lines), varying only the level of prediction (*α*_*P*_, blue), and varying only the level of series resistance compensation (*α*_*R*_, red), showing distinctive effects between the two when applied to *I*_Na_.

First, the different responses to the prediction levels (*α*_*P*_, blue) were examined without series resistance compensation. The current was left-shifted (peaks occurred earlier) as the level of prediction/supercharging compensation increased; the phenomenon was captured by the new voltage-clamp model, and the model was even able to capture the small hump at the beginning of the voltage step, demonstrating the power of the computational model. This effect can be explained by looking at the simulated membrane voltage (top panel) which is a consequence of ‘loss of voltage clamp’, and as the prediction was applied, it pulled the membrane voltage together with the measured current back with a smaller delay of the peak.

Next, the responses to the series resistance compensation were compared (*α*_*R*_, red) with *α*_*P*_ = 80 %. In this case, instead of correcting the delay of the current, series resistance compensation improved the amplitude of the measured current by bringing the membrane voltage closer to the command voltage. At the highest possible compensation level, *α*_*R*_ = *α*_*P*_ = 80 %, the measured current (dark red) was much closer to the ideal current (black dashed line). However, even that was not enough to obtain the desired voltage step (as shown by the simulation) which can result in erroneous estimates of the underlying physiological characteristics, as demonstrated in the next section. Repeats of the experiments are shown in the **Supplementary Materials**.

### 2.4 Extrapolating to physiological I-V relationships with best available (imperfect) data

We further demonstrate a new way of using our computational model of voltage-clamp experiments for extrapolating current-voltage (I-V) relationships of imperfect data (even under the best experimentally-possible conditions, they are still imperfect) to the potential underlying physiological I-V relationships for corrected maximum current conductance estimation. Unlikely the previous section, where strong experimental artefact was desired for assessing the computational model performance, here, the best experimentally-possible setup closest to physiological conditions was employed. The *in vitro* experiments were performed with HEK cells over-expressing Na_V_1.5 under rupture-patch conditions, in order to achieve lowest possible series resistance in the experiments (see **Methods** for details).

Fitting a mathematical model of fast sodium current (Iyer et al., 2004) with the voltage-clamp model, with changing only the maximum current conductance, *R*_s_, *C*_m_, and 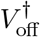, to the 80 % compensation (the highest possible level of compensation in practice) experimental data allows us to elucidate the effects of compensations (shown in different colours) and to predict the artefact-free, true physiological condition (dotted lines), as shown in Fig. 5. The effects of compensations are distinctive on I-V relationships, which is evidence from the experimental data (solid lines with crosses; Fig. 5B) and is confirmed by the voltage-clamp model (dashed lines with dots; Fig. 5A); the effects can be explained by the model as described later in this section. Increasing the compensation level not only elevated the maximum current but also right-shifted the peak of the I-V curves. More importantly, the predicted artefact-free condition I-V curve was also different to any of the (imperfectly) compensated conditions. Therefore, our computational approach provides a gateway to understanding the true underlying physiological I-V relationships of Na_V_1.5 and the corrected maximum current conductance estimation that otherwise cannot be measured experimentally.

**Figure 5:**
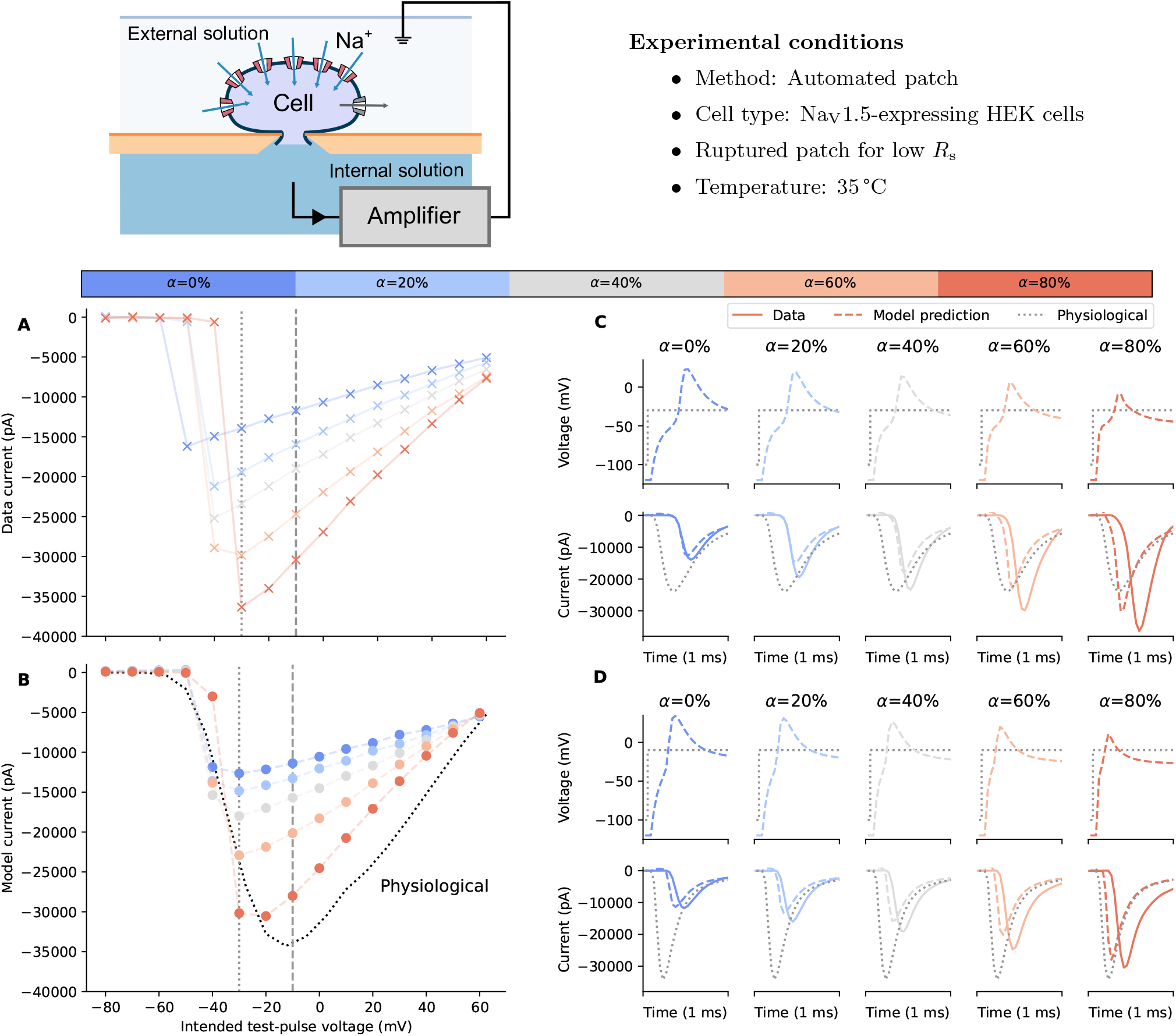
Correction of fast sodium current experiments using the voltage-clamp experimental artefact model. **Top:** A schematic of the experimental setup and the experimental conditions for the ruptured patch experiments with HEK cells over-expressing Na_V_1.5. **Bottom: (A)** Experimental recordings (solid lines) of the current-voltage (I-V) relationships of fast sodium current, measured with different levels of series resistance and prediction compensation (*α*). **(B)** Simulated (dashed lines) of the I-V relationships. **(C, D)** Elicited current and membrane voltage of two voltage steps indicated on the I-V curves (A, B). Left to right show effects of different compensation levels (*α*), with the experimental recording shown in solid lines and simulated effects shown in dashed lines. In all panels, the corrected fast sodium current is shown in black dotted lines.

Fig. 5C-D shows the current and membrane voltage of two of the voltage steps where the experimental current peaked (*−*30 mV) and the artefact-free condition peak (*−*10 mV), which demonstrates the causes of the I-V relationship contaminations under imperfect compensation. The differences caused by different compensation levels can be explained by visualising the loss of voltage clamp, i.e. the cell experienced different voltages when clamped with different compensation levels, resulting in different current dynamics. More repeats of the *in vitro* experiments are shown in **Supplementary Material**. Although we note that typical experiments would not vary the compensation levels during the experiment, this new type of experiments allowed us to verify the effectiveness of the voltage-clamp model and demonstrate the ‘trend’ of the compensation levels.

### 2.5 Averaging I-V curves does not eliminate experimental artefacts

Finally, we found that conventional averaging of I-V curves may obscure the problems of imperfect compensation, which may deceive us into thinking it was a good approximation to the physiological conditions. Fig. 6 shows the results of taking averages of simulated I-V curves (blue; averaged from transparent grey lines) compared to the underlying physiological model I-V curve (red). Each individual I-V curve was computed by simulating the voltage-clamp model at 80 % compensation level with the O’Hara et al. (2011) fast sodium current, where the scaling of the current maximum conductance, membrane capacitance, and series resistance were sampled through a Latin hypercube algorithm with the boundaries [0.2, 5], [4, 15] MΩ, and [8, 22] pF, respectively. The capacitance and series resistance were assumed to be estimated perfectly, so 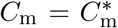 and 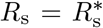 Simply taking averages of 80 % compensation (typical experimental compensation level) I-V curves produced a smooth I-V curve. However, neither the peak voltage / current nor the gradient of the average I-V curve (blue) match those of the underlying physiological I-V curve (red), therefore the average I-V does not hold as a good approximation of the physiological I-V (Fig. 6). A typical sample size *n* = 5 was first tested, and subsequently we increased number of samples to *n* = 25 and *n* = 75. In all cases the maximum peak current was underestimated by 10 – 30 %, the peak voltage was left shifted by about 10 – 25 mV, and the gradient of the I-V were different. Increasing the number of samples and taking averages does not approximate the underlying physiological I-V curve well. Also note that the experimental ‘error bar’ limits would not be close to the underlying physiological properties either, even if standard deviations rather than standard errors of the mean (SEM) were to be shown (see **Supplementary Material**). Normalising I-V curves before averaging would not rescue the issue either, as shown in **Supplementary Material**.

**Figure 6:**
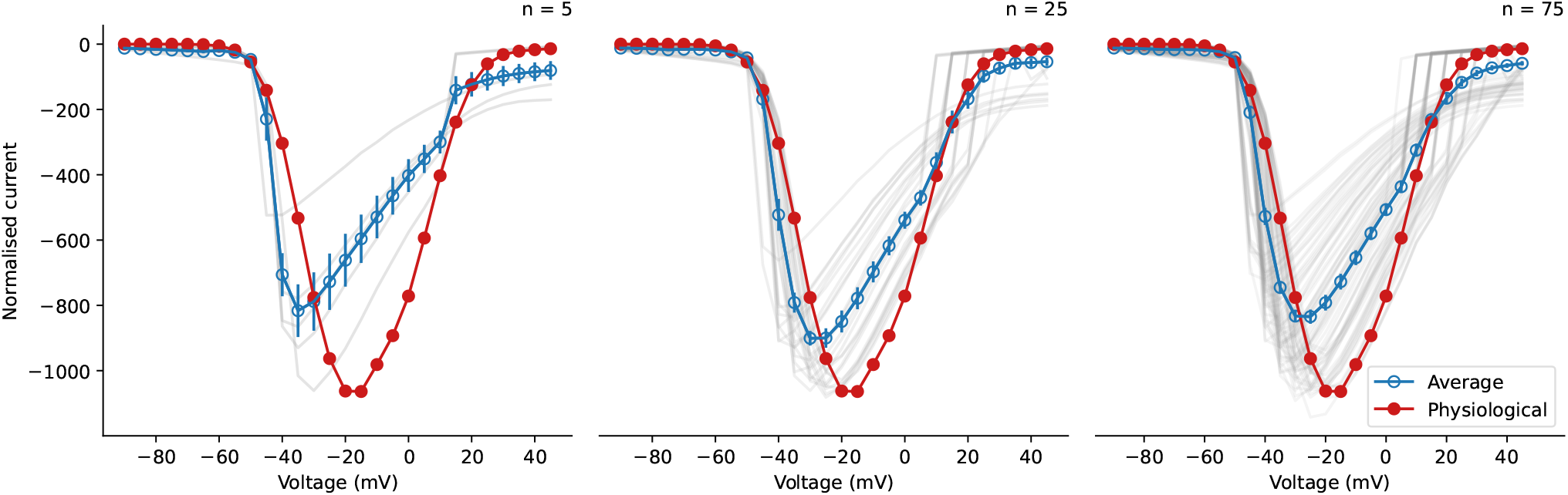
Consequences of averaging fast sodium current I-V curves of multiple runs of experiments. Comparing the average simulated I-V curve (blue with error bars showing SEM; averaged from transparent grey lines) to the artefact-free physiological model I-V curve (red), neither the peak current, the peak voltage, nor the gradient of the curve of the average I-V matches the physiological I-V curve. Left to right shows different number of repeats, *n* = 5, 25, and 75, respectively.

### 2.6 Experimental artefacts may lead to mischaracterisation of mutants or drug effects

To further illustrate the potential impact of the experimental artefacts, we performed a synthetic experiment here: Consider a Na_V_1.5 mutant or a compound that blocks Na_V_1.5, which its true physiological effect is reduction in current conductance (e.g. via trafficking inhibition or simple pore block), which is characterised via patch-clamp voltage-clamp experiments and compared against the wild type (WT) or control in the same experimental settings. As an illustrative example (see e.g. Amin et al., 2005), we considered the case which the effect of the mutant or drug block (orange) is a reduction to one-third of the current conductance compared to the WT or control (blue), as illustrated in Fig. 7 (left). To mimic the real experimental situations, we sampled the experimental conditions following *R*_s_ *∼* LogNormal(2.5, 1.5) MΩ, *C*_m_ *∼* LogNormal(40, 10) pF, and 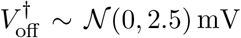, where LogNormal(*μ, σ*) and *𝒩* (*μ, σ*) are a log-normal distribution and a normal distribution with mean *μ* and variance *σ*, respectively, while each virtual cell has the same current density.

**Figure 7:**
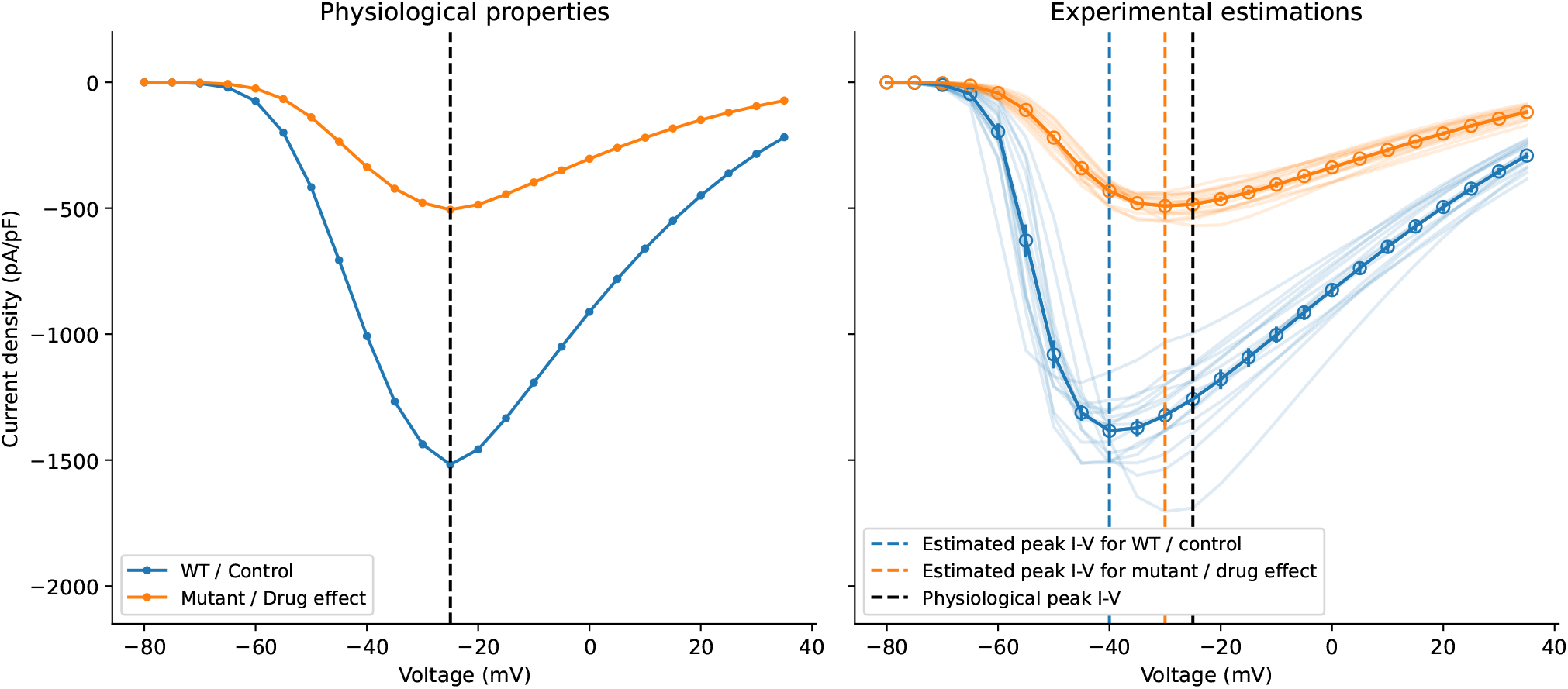
An example of how experimental artefacts may lead to mischaracterisation of mutants or drug effects. **(Left)** The true underlying physiological I-V curves of the mutant or the drug (orange) compared to the WT or control (blue), where only current conductance was reduced by one-third. **(Right)** The observed I-V curves of the two conditions under one of the most stringent experimental conditions (only *R*_s_ *<* 4 MΩ was allowed). The solid lines are the mean of the individual I-V curves (*n* = 15) shown in transparent lines, with error bars showing the SEM. Vertical dashed lines show the voltage at which I-V curves peak, with blue for WT/control, orange for mutant/drug, and black for the true effect.

Even when following one of the most stringent experimental conditions in the literature, where only *R*_s_ *<* 4 MΩ were accepted through quality controls (for details of the settings, see **Methods**), we found that the recorded I-V relationships (*n* = 15) were deceiving and would lead us to interpret as a mutant or a drug that “not only reduces the current magnitude *but also* shifts the I-V curves by 10 mV”, as shown in Fig. 7 (right). Again, similar to the previous section, the experimental ‘error bar’ (SEM) limits would not be close to the underlying physiological properties. The consequences will get even worse if the quality control is less stringent. The results strongly suggest the importance of this work that presents a new computational approach for handling experimental artefacts in patch-clamp voltage clamps.

## 3 Discussion

Patch-clamp voltage clamp is the gold standard for cellular electrophysiology. Yet, the experimental error which is shown to be persistent even after typical compensations has not been properly addressed, leading to the pervasiveness of (ignored) artefacts in the literature. Here, such effects of artefact have been addressed and explained through a new, validated computational model of voltage-clamp experiments. By elucidating the forms of these artefacts using the fast sodium current as an example, it has been further shown that averaging of data in a conventional manner does not eliminate the effects of artefacts — they leave a systematic bias. The results reveal not only how artefacts affect the experimental recordings, but also emphasise the importance of addressing these effects, as we illustrated how artefacts may confound our characterisation of mutant or drug effects.

Typically, one might attempt to reduce the effects of the artefacts through adjusting to unphysiological conditions in the experiments, e.g., by recording at lower temperatures, using unphysiological solutions to reduce current flow, etc. However, we have previously observed how extrapolating from unphysiological temperatures to physiological ones can produce biased results (Lei et al., 2019), and similar effects may occur if channel kinetics or maximal conductances are sensitive to ion concentrations as well. We argue that our computational models and simulations offer a new way to use heavily-polluted data, making the study of physiological conditions in experiments possible, which is vital for studying ion channel electrophysiology without extrapolation introducing further bias.

The computational model of the patch-clamp voltage clamp has been fully validated through a novel design of an electrical model cell circuit. Unlike biological cells, in which we do not have a full understanding of physiological electrical properties, a model cell made of physical electrical components is fully transparent. The electrical model cell was designed to create a fast response current, to allow a controlled experimental validation for new components of our computational model that were not included in Lei et al. (2020a). This validation allows us to separate our uncertainty in ion channel physiology and patch-clamp voltage clamp, as we are interested in the latter here. Our results (Fig. 3) show that, with the correct model of the electrical model cell, we are able to reproduce the effects of various levels of machine compensation, reassuring us as to the robustness of the model.

In our analysis of the voltage-clamp model subject to different compensation settings with a fast sodium current model, we observed independent effects of series resistance compensation and prediction, as further confirmed by experimental data from fast sodium current (Fig. 4). Although one would not vary or use compensation levels (*α*) less than 70 % (but typically not higher than 85 %) when studying the physiological properties of the cells, our results reproduce the trend of the compensation effects, providing a qualitative indication of the likely over/under estimation of recorded currents through time. Similarly, Fig. 5 also shows a similar effect when studying I-V relationships as typically done in the literature. The observed shifts and biases were due to the loss of control of the membrane voltage that the cell experienced, and we expect other derived biomarkers, such as observed reversal potential, would also be shifted due the change in the membrane voltage. These effects are systematic biases of the experiments (Fig. 6) which cannot be eliminated by typical averaging of multiple repeats of the same experiment. Therefore, in voltage-clamp experiment, the experimental artefacts will lead to incorrect estimations of the cell properties.

Fig. 4 also shows that not only the I-V relationships but also the time-to-peak would be affected by the artefacts. However, such effects on the time-to-peak are neither linear nor straightforward, in fact, when close to the activation threshold voltage, the artefact effects are seemingly unpredictable. Fig. 8A shows the experimental recordings of the fast sodium current at different compensation levels (left to right). The red arrows indicate the recorded delays of the peak currents at around *−*30 to *−*40 mV. Depending on the conditions, the delay can be longer than the time taken for inactivation, see for example *α* = 40 %. Even for the typical experimental conditions, with *α* = 80 %, we can still observe weaker but significant delays in some of the peaks. These phenomena can be explained using our computational models, as shown in Fig. 8B. Using the sodium model from Gray and Franz (2020) together with our voltage-clamp mathematical model (middle panel), we are able to qualitatively reproduce the seemingly unpredictable delay patterns (experimental data in the left panel). Our model reveals that the delays were due to the nonlinear interactions between the loss-of-clamp of the membrane voltage and the sudden large current activated near the threshold voltage, as shown in the middle panel of Fig. 8B.

**Figure 8:**
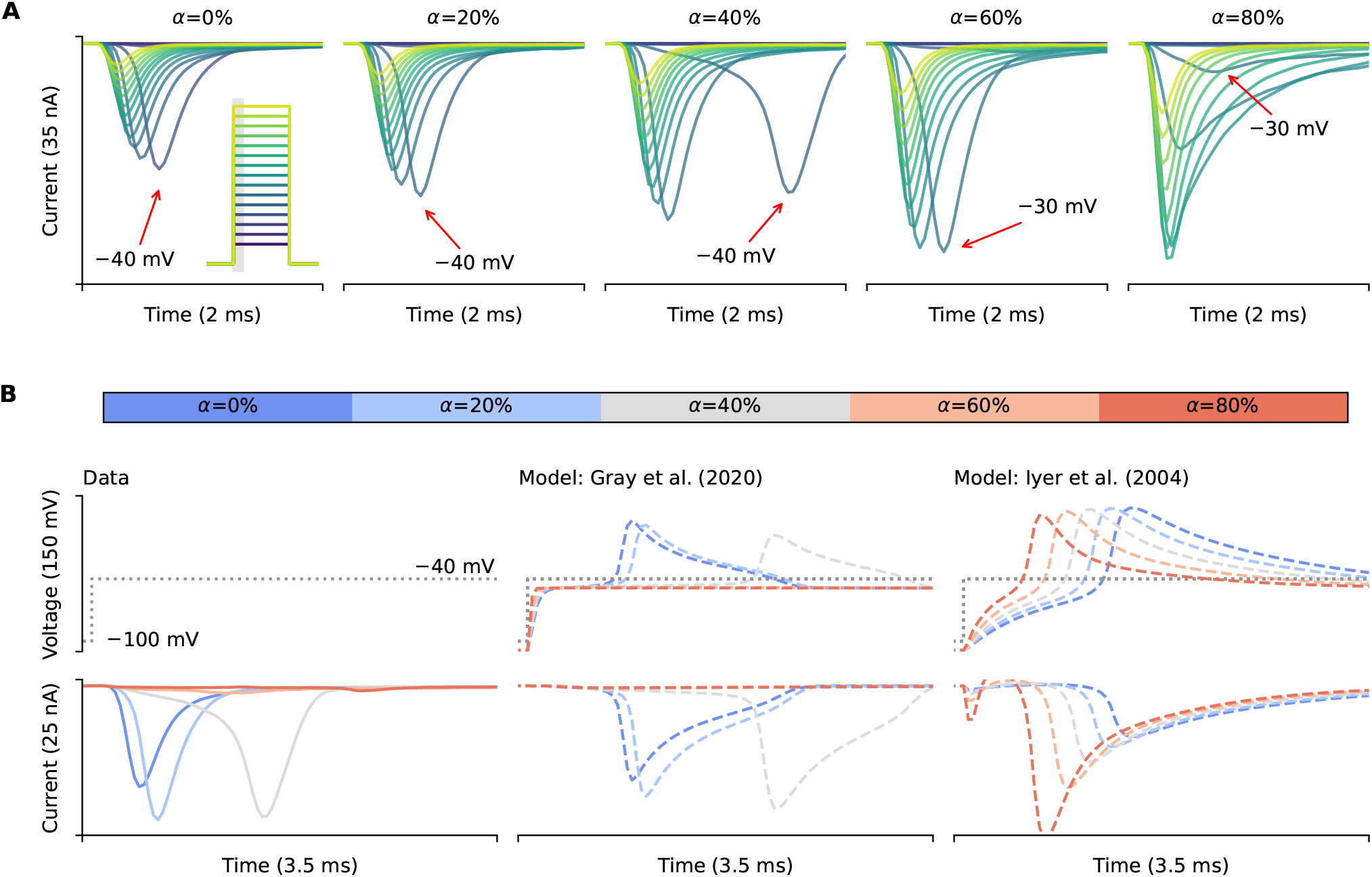
The voltage-clamp model can qualitatively capture the delayed current of fast sodium current. **(A)** Experimental recordings of fast sodium current activation at different levels of compensation. It shows that near the threshold voltage of current activation (*∼−*30 to *−*40 mV), there are delays in the peak time of the current as indicated by red arrows. Inset, the voltage protocol, with holding potential at *−*100 mV. **(B)** An examination of the *−*40 mV test step. From left to right are experimental data (true membrane voltage is unknown) and voltage-clamp model simulations with Gray and Franz (2020) and Iyer et al. (2004), respectively, with different levels of compensation (*α*).

A potential explanation to the seemingly unpredictable time-to-peak delay was due to a positive feedback between the current and the loss of clamp in membrane voltage. Due to the nature of a very steep activation I-V relationship of the sodium current, when the voltage is set to just below the activation threshold voltage of the current, a small negative current from the cell would induce a small positive voltage due to the series resistance effect. This induction further induces a slightly bigger sodium current to push the membrane voltage even closer to the activation threshold. This positive feedback will eventually set the membrane voltage to hit the activation threshold of the sodium current, which results in the large current spike with a delay, therefore this time-to-peak delay will depend on the series resistance and its compensation, as well as the slope of the sodium I-V activation curve.

However, such phenomena are sensitive to the correctness of the current activation in the physiological model. Although Gray and Franz (2020) model simulations qualitatively match the experimental delays due to the loss-of-clamp of the membrane voltage, this behaviour is sensitive to the choice of model. We were not able to reproduce the observed delay patterns by simply replacing the ionic current model with the fast sodium current from Iyer et al. (2004), as shown in the right panel of Fig. 8B. We believe this is due to high accuracy required in model current-voltage relationships near *−*30 to *−*40 mV to reproduce the delay patterns. As discussed above, we note that a limitation of the approach, shared by Lei et al. (2020a); Montnach et al. (2021); Abrasheva et al. (2024), is the reliance on the assumption of a known ion channel model. Any discrepancy between the mathematical ion channel model and the real currents may cause nonlinear disruption as demonstrated here.

As a final remark, we note that the voltage-clamp model captures the first-order behaviour of the amplifier components, as some of the timescales of the interactions between amplifier electrical components are in the order of microseconds or in a very high frequency domain which are not essential for correcting the experimental errors and biases for most of the systems of interest. For example, some of the amplifiers use a specific type of low-pass filter, such as a 4-pole low-pass Bessel filter on the HEKA EPC 10 amplifier, for Eq. (9). There may also be other terms such as nonlinear leak currents in some automated patch clamp (Lei et al., 2020b) that are not included in our model that may be setup or experiment specific. Therefore, Eqs. (1)–(10) form a first-order approximation to the overall effect which should be sufficient for capturing and correcting the majority of experimental errors, as demonstrated here, when experiments are performed according to manufacturers’ recommended procedures.

In conclusion, our new modelling approach provides a step forward in how voltage-clamp electrophysiological data can be interpreted, making fitting, and thus recovering, accurate estimates of biophysical properties possible.

## 4 Methods

### 4.1 Derivation of the mathematical compensation model

There are two effects introduced by *R*_s_, the first one causes *V*_m_ to deviate from *V*_cmd_ and the second slows down *V*_m_’s approach to *V*_cmd_. The first effect is caused by (*I*_ion_ + *I*_leak_), which can be reduced through a series resistance compensation (Sigworth et al., 1995; Sigworth, 1995a; Weerakoon et al., 2009). The series resistance compensation therefore sets the voltage clamp to 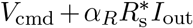 instead of *V*_cmd_, where 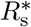 sis the machine estimation of the series resistance R_S_, and *α*_*R*_ is the requested proportion of series resistance compensation (a machine setting, typically 70–85 %).

The second effect is caused by the product of the series resistance *R*_s_ and the membrane capacitance *C*_m_, i.e. the membrane assess time constant, which can be reduced through a compensation termed “super-charging” (Sigworth, 1995b) by clamping voltage to a large overshoot (hence the name “supercharging”) proportional to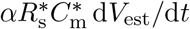. Therefore, we have

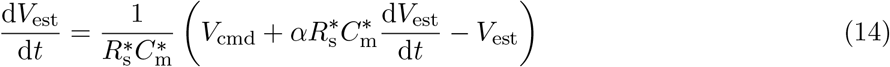

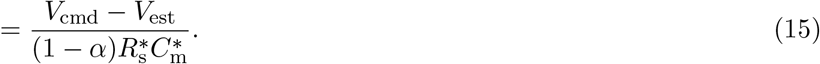

The effect of the overshooting is to charge the membrane capacitance quickly. We note that supercharging is particularly important when measuring big, very fast currents such as *I*_Na_. The full derivation of the model is included in **Supplementary Material**.

### 4.2 Electrical model cell validation experiments

The electric circuit of the model cell was soldered with the electrical components listed in Fig. 3 (top). The two probes were each connected to a multichannel patch-clamp amplifier (HEKA EPC 10 USB multichannel version), one was set to voltage-clamp mode and the other was in current-clamp mode (with zero current injection), as shown in Fig. 3. Automated fast capacitance estimation (i.e.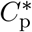) was applied using the HEKA PATCHMASTER software to compensate for *C*_p_, while the values of 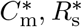 were set to the model cell values during the experiments. Three types of compensation settings were tested: (1) the supercharging (prediction, *α*_*P*_) was fixed to zero and series resistance compensation (*α*_*R*_) was varied from 0 % (no compensation) to 80 % with an increment of 20 %, and 95 %; (2) *α*_*R*_ was set to the maximum (95 %) and *α*_*P*_ varied from 0 % to its maximum; and (3) *α*_*R*_ and *α*_*P*_ were varied together. Experimental data were acquired with a large voltage step protocol that consists of a holding step *−*80 mV stepping to 50 mV for 20 ms before stepping back to the holding step. The protocol was intended to draw a large current response from the electrical model cell.

### 4.3 hiPSC-CM perforated patch experiments

The perforated patch-clamp experiments with hiPSC-CMs (Fig. 4) are from a subset of cells we included in a recent publication (Clark et al., 2022) — in the next couple of paragraphs, we provide a brief overview of the experiments. The data included in this manuscript (i.e., Na_V_ protocols), however, have not been previously published. The data reported here were collected after the acquisition of the current clamp and voltage clamp data published in Clark et al. (2022).

Briefly, frozen stocks of hiPSC-CMs from a healthy individual (SCVI-480CM) were obtained from Joseph C. Wu, MD, PhD, at the Stanford Cardiovascular Institute Biobank. The hiPSC-CM line was derived from an African-American female donor and was approved by Stanford University Human Subjects Research Institutional Review Board. We used Amphotericin B as the perforating agent. Patch-clamp measurements were made at 37 °C and at 10 kHz by a patch-clamp amplifier (Model 2400; A-M Systems, Sequim, WA) controlled by the Real Time eXperiment Interface (RTXI; http://rtxi.org) to send commands to the amplifier via the data acquisition card (PCI-6025E; National Instruments, Austin, TX). *R*_m_, *C*_m_, and *R*_s_ values were measured at 0 mV within one minute prior to the acquisition of Na_V_ protocol data.

We acquired I-V curves by stepping from *−*80 mV to voltages between *−*70 mV and 60 mV, incrementing by 10 mV. We held the excitation step for 50 ms before returning to *−*80 mV for 400 ms. To test multiple compensation settings, we first set series resistance compensation (i.e., *α*_*R*_) to 0 %, and increased supercharging (i.e., *α*_*P*_) from 0 % to 80 % in 20 % increments. Then, we set supercharging to 70 % and incrementally increase series resistance compensation from 0 % to 80 % in 20 % increments. Compensation settings above the maximum here (*α*_*P*_ = 70 %, and *α*_*R*_ = 80 %) resulted in undesirable oscillations.

### 4.4 Na_V_1.5 experiments

Na_V_1.5-expressing HEK cells were purchased from Charles River. Experiments were conducted at 35 °C using a four-channel Nanion Patchliner and either medium or high resistance chips. Series resistance compensation and supercharging were initially set at 0 % and incremented by 20 % up to 80 %. The I-V recordings were collected by holding at *−*100 mV for 1980 ms before stepping to an incrementally increasing voltage between *−*80 mV and 60 mV and holding for 20 ms. The internal solution included: 10 mM EGTA, 10 mM Hepes, 10 mM CsCl, 10 mM NaCl, 110 mM CsF, and with a pH adjusted to 7.2 with CsOH (*>* 280 mOsm). The external solution included: 140 mM NaCl, 4 mM KCl, 2 mM CaCl_2_, 1 mM MgCl_2_, 5 mM D-Glucose monohydrate, 10 mM Hepes, and with a pH adjusted to 7.4 with NaOH (298 mOsm).

### 4.5 Fitting to experimental data

Model prediction results in Fig. 5 were obtained by fitting the Iyer et al. (2004) fast sodium model with the voltage-clamp model to the experimental data. Only the maximum conductance of the current and the voltage-clamp model parameters were varied and fitted to the data, where the voltage-clamp model parameters include *R*_s_, *C*_m_, 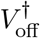, and *g*_leak_. The other parameters in the voltage-clamp model, namely 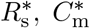, and 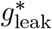 were set to the amplifier provided estimated. Four of the fitted parameters *p*_*i*_: maximum conductance, *R*_s_, *C*_m_, and *g*_leak_ were transformed to *s*_*i*_ during the optimisation using 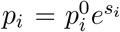, where 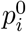 were the reference values of the parameters which were 600 nS, 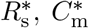, and 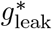, respectively. The optimisation was repeated 40 times at random initialisation of the starting points, to ensure obtaining the global optimum solution, using the covariance matrix adaptation evolution strategy (CMA-ES) implemented in Clerx et al. (2019).

### 4.6 Simulations of current-voltage replication

Simulations in Fig. 6 were generated using the O’Hara et al. (2011) sodium current model with the voltage-clamp artefact model included. I-V curves were generated by holding at *−*100 mV for 2000 ms before stepping to an incrementally increasing voltage between *−*90 and 50 mV and holding for 20 ms at each step. Both the supercharging and series resistance compensation parameters were set to 80 % for all simulations, while sodium conductance, series resistance, and capacitance was varied. Latin hypercube sampling was used to select the parameters for all models. The sodium conductances were selected from a log-transformed scale between 0.2*×* and 5*×* the baseline values. All capacitances were between 4 MΩ and 15 MΩ and series resistances between 8 and 22 pF. We assumed the capacitance and series resistance were estimated perfectly, so 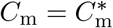 and 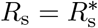.

### 4.7 Simulations of mutations or drug effects

Simulations in Fig. 7 were generated using the Iyer et al. (2004) sodium current model together with the voltage-clamp model. The mutant or drug effect in this simulation study was assumed to have the exact same physiological properties of the WT or control condition (the original model setting) except the maximum current density was assumed to be one third of the WT/control condition. I-V curves were generated with the same protocol as in Section 4.6. The maximum current density was set to be 1500 A*/*F and 500 A*/*F for WT/control and mutant/drug effect, respectively, for the I-V curves under ideal voltage clamp. During the experimental estimation using the voltage-clamp model, the supercharging and series resistance compensation parameters were set to 80 % for all simulations. Only *R*_s_, *C*_m_, and 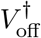 were varied during each realisation of the experiment as in the reality. To ensure the quality of the experiment realisation, the parameters were sampled following *R*_s_ *∼* LogNormal(2.5, 1.5) MΩ, *C*_m_ *∼* LogNormal(40, 10) pF, and 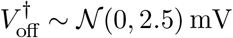 for both WT/control and mutant/drug conditions, where LogNormal(*μ, σ*) and 𝒪 (*μ, σ*) are a log-normal distribution and a normal distribution with mean *μ* and variance *σ*^2^, respectively. The maximum conductance of the current was scaled according to the sampled *C*_m_ to give the same current density (under the ideal voltage clamp). We further assumed a reasonably small 5 % estimation error of the series resistance and membrane capacitance, which was done by sampling 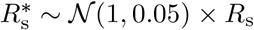 and 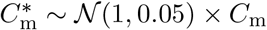

## Data Availability

All code and data are available at https://github.com/CardiacModelling/nav-artefact-model.

## Acknowledgments

This work was performed in part at the High Performance Computing Cluster supported by the Information and Communication Technology Office of the University of Macau.

