## Supplementary Material for "Resolving artefacts in voltage-clamp experiments with computational modelling: an application to fast sodium current recordings"

***Author affiliations as they appear in the main article***

#### S1 Model derivation

The derivation of the equations governing a voltage-clamp experiment *without* any compensation follows exactly as Lei et al. (2020), which is not repeated here. These include the equations for the effects of membrane capacitance, series resistance, leak current (seal resistance), pipette capacitance, and amplifier delays. Below shows the derivation of the mathematical model of how modern patch amplifiers typically compensate for them (Axon Instruments Inc., 1999; HEKA Elektronik GmbH, 2018), following the derivation from (Lei, 2020). Firstly, the voltage offset is usually estimated and compensated *prior* to adding the cell to the system, either with an automated correction estimated using *software control* or by applying manually a voltage offset such that it gives zero current when clamped at zero voltage, so the compensation circuit is not shown in our patch clamp equivalent circuits (Neher, 1995; Sigworth et al., 1995). The major source of voltage offset may be the liquid junction potential, a potential difference of $\sim 2 - 12\,\mathrm{mV}$ which develops when the pipette-filling solution is different from the bath solution (Neher, 1992). The adjustment is usually done by adding the theoretically estimated liquid junction potential offset to $V^*_{\mathrm{off}}$. We can write the error in the estimate of the overall voltage offset $V^\dagger_{\mathrm{off}}$ as

$$V^\dagger_{\mathrm{off}} = V_{\mathrm{off}} - V^*_{\mathrm{off}}. \tag{S1.1}$$

We then simply need to replace all instances of $V_{\mathrm{off}}$ in the equations above with $V^\dagger_{\mathrm{off}}$ to describe the effect of imperfect voltage offset compensation, and $V^\dagger_{\mathrm{off}}$ is assumed to be $\mathcal{O}(10)\,\mathrm{mV}$.

Secondly, to compensate the effect of the parasitic capacitance at the electrode, an additional current is injected at the electrode to compensate for the current drawn by the parasitic capacitance. By analysing the fast capacitance compensation part, we obtain the compensation current as $C^*_{\mathrm{p}}\frac{\mathrm{d}V_{\mathrm{clamp}}}{\mathrm{d}t}$ where $C^*_{\mathrm{p}}$ is the amplifier's estimate of the parasitic capacitance $C_{\mathrm{p}}$. Then we have

$$I_{\mathrm{m}} = I_{\mathrm{in}} + C^*_{\mathrm{p}}\frac{\mathrm{d}V_{\mathrm{clamp}}}{\mathrm{d}t} - I_{\mathrm{p}} - I_{\mathrm{leak}} - I_{\mathrm{ion}}, \tag{S1.2}$$

and

$$I_{\mathrm{in}} = I_{\mathrm{ion}} + I_{\mathrm{leak}} + C_{\mathrm{m}}\frac{\mathrm{d}V_{\mathrm{m}}}{\mathrm{d}t} + \left(C_{\mathrm{p}}\frac{\mathrm{d}V_{\mathrm{p}}}{\mathrm{d}t} - C^*_{\mathrm{p}}\frac{\mathrm{d}V_{\mathrm{clamp}}}{\mathrm{d}t}\right). \tag{S1.3}$$

This is usually known as 'C-Fast' compensation.

Thirdly, we need to consider compensation for the cell membrane capacitance $C_{\mathrm{m}}$. Usually the effect of $C_{\mathrm{m}}$ is reduced by a hardware 'C-Slow' compensation, using a similar circuit to the 'C-Fast' compensation discussed above (Sigworth et al., 1995; Sigworth, 1995a). However, since the value of $C_{\mathrm{m}}$ can reach 100 pF in some cell types, and capacitor sizes can be limited, the 'C-Slow' compensation is sometimes performed as a post-processing step by the amplifier control software rather than using built-in amplifier hardware (Weerakoon et al., 2010). In either case, the full capacitance compensation can be written as

$$I_{\mathrm{in}} = I_{\mathrm{ion}} + I_{\mathrm{leak}} + \left(C_{\mathrm{m}}\frac{\mathrm{d}V_{\mathrm{m}}}{\mathrm{d}t} - C^*_{\mathrm{m}}\frac{\mathrm{d}V_{\mathrm{est}}}{\mathrm{d}t}\right) + \left(C_{\mathrm{p}}\frac{\mathrm{d}V_{\mathrm{p}}}{\mathrm{d}t} - C^*_{\mathrm{p}}\frac{\mathrm{d}V_{\mathrm{clamp}}}{\mathrm{d}t}\right), \tag{S1.4}$$

where $C_\mathrm{m}^*$ is the amplifier (or user's) estimate of the membrane capacitance $C_\mathrm{m}$, and $V_\mathrm{est}$ is given below.

Finally, in voltage clamp, we want $V_\mathrm{m}$ to approach $V_\mathrm{cmd}$ as quickly as possible. However, there are two effects introduced by $R_\mathrm{s}$, the first one causes $V_\mathrm{m}$ to deviate from $V_\mathrm{cmd}$ and the second slows down $V_\mathrm{m}$'s approach to $V_\mathrm{cmd}$. The first effect is caused by $(I_\mathrm{ion} + I_\mathrm{leak})$, which can be reduced through a series resistance compensation (Sigworth et al., 1995; Sigworth, 1995a; Weerakoon et al., 2009). By analysing the series resistance compensation part, instead of clamping to $V_\mathrm{cmd}$, it is set to $V_\mathrm{cmd} + \alpha R_\mathrm{s}^* I_\mathrm{out}$, where $R_\mathrm{s}^*$ is the machine estimation of the series resistance $R_\mathrm{s}$, and $\alpha$ is the requested proportion of series resistance compensation (a machine setting, typically 70–85 %).

The second effect is caused by the product of the series resistance $R_\mathrm{s}$ and the membrane capacitance $C_\mathrm{m}$, i.e. the membrane assess time constant $\tau_\mathrm{a}$, which can be reduced through a compensation termed "supercharging" (Sigworth, 1995b). We set the clamping voltage to have a large overshoot (hence the name "supercharging") proportional to $\alpha R_\mathrm{s}^* C_\mathrm{m}^* \,\mathrm{d}V_\mathrm{est}/\mathrm{d}t$, where

$$\frac{\mathrm{d}V_\mathrm{est}}{\mathrm{d}t} = \frac{1}{R_\mathrm{s}^* C_\mathrm{m}^*}\left(V_\mathrm{cmd} + \alpha R_\mathrm{s}^* C_\mathrm{m}^* \frac{\mathrm{d}V_\mathrm{est}}{\mathrm{d}t} - V_\mathrm{est}\right) \tag{S1.5}$$

$$= \frac{V_\mathrm{cmd} - V_\mathrm{est}}{(1-\alpha) R_\mathrm{s}^* C_\mathrm{m}^*}, \tag{S1.6}$$

according to Sigworth (1995b, Figure 18). The effect of the overshooting is to charge the membrane capacitance quickly. Including all the compensations, $V_\mathrm{clamp}$ becomes

$$\frac{\mathrm{d}V_\mathrm{clamp}}{\mathrm{d}t} = \frac{1}{\tau_\mathrm{sum}}\left(\left(V_\mathrm{cmd} + \alpha R_\mathrm{s}^*\left(I_\mathrm{out} + C_\mathrm{m}^* \frac{\mathrm{d}V_\mathrm{est}}{\mathrm{d}t}\right)\right) - V_\mathrm{clamp}\right), \tag{S1.7}$$

to counterbalance the two effects caused by the series resistance. Note that the supercharging correction is particularly important when measuring big, very fast currents such as $I_\mathrm{Na}$ which has a time-to-peak within ∼5 ms, however this correction poses almost no issue when analysing smaller, slower currents, for example $I_\mathrm{Kr}$.

### Supplementary Figures

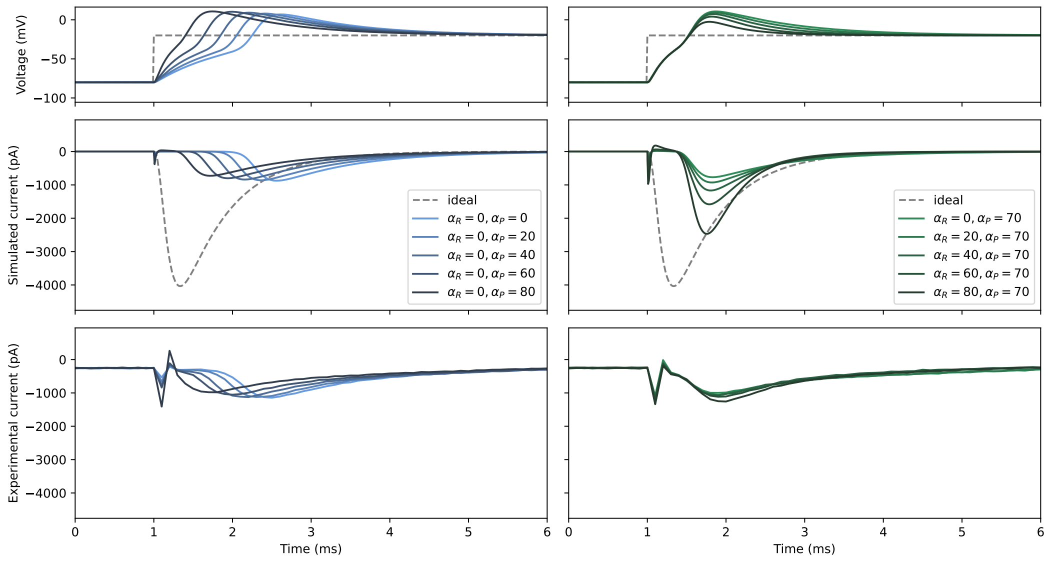


Figure S1: Forward sensitivity analysis of the experimental artefact of fast sodium current using Gray and Franz (2020) and hiPSC-CMs (cell 0).

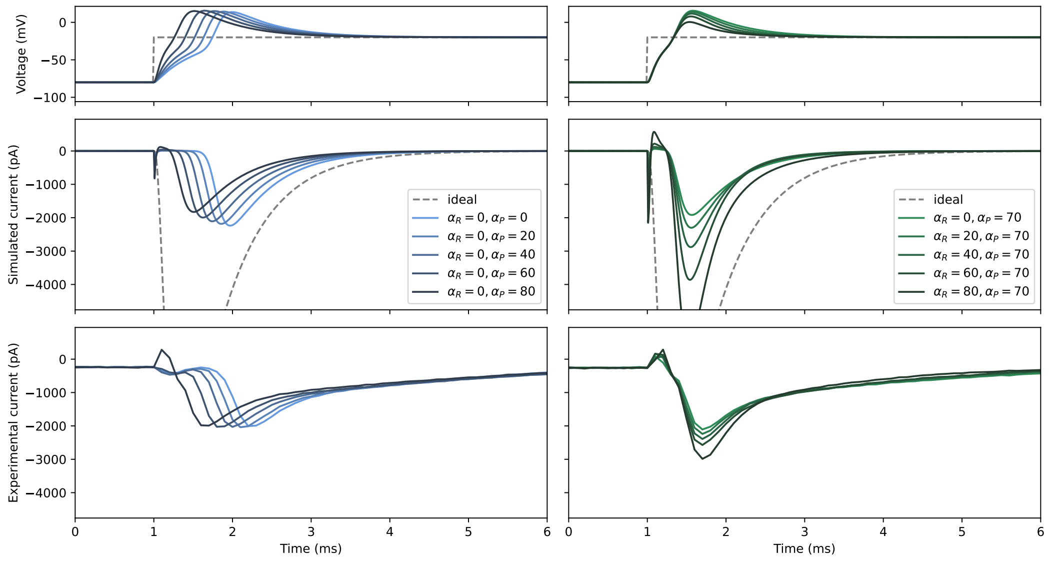


Figure S2: Forward sensitivity analysis of the experimental artefact of fast sodium current using Gray and Franz (2020) and hiPSC-CMs (cell 1; same cell as in the main text figure).

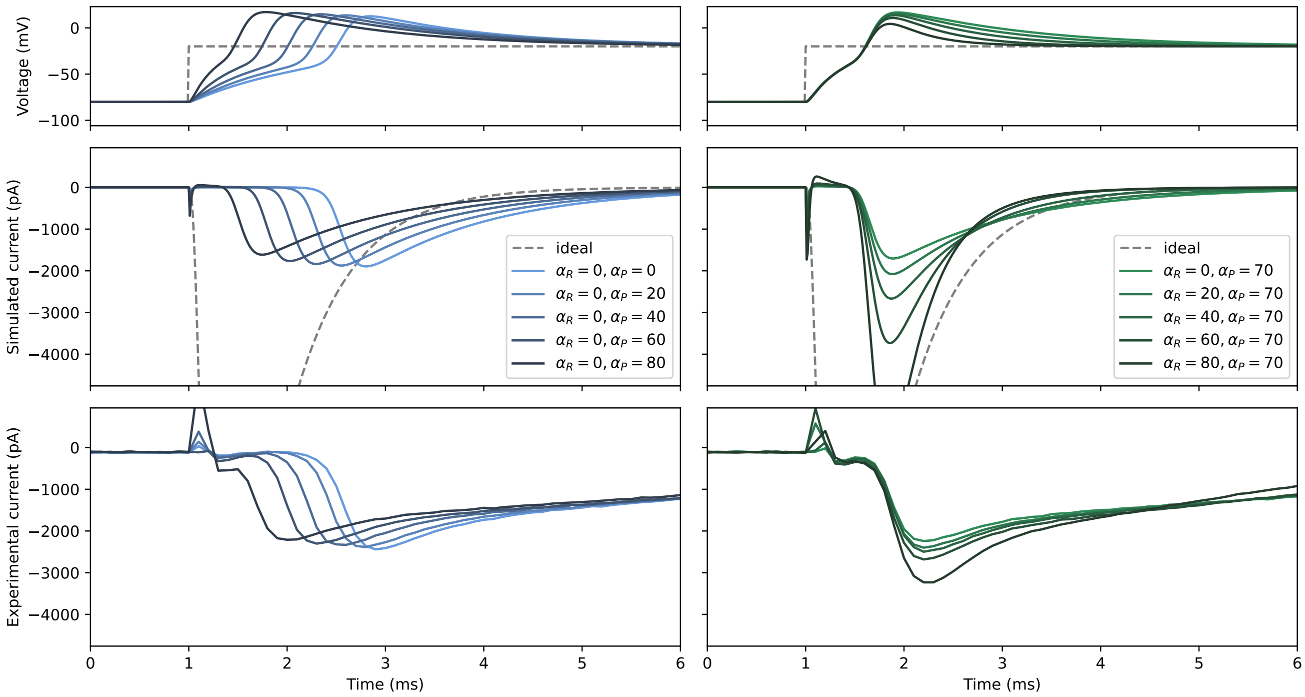


Figure S3: Forward sensitivity analysis of the experimental artefact of fast sodium current using Gray and Franz (2020) and hiPSC-CMs (cell 2).

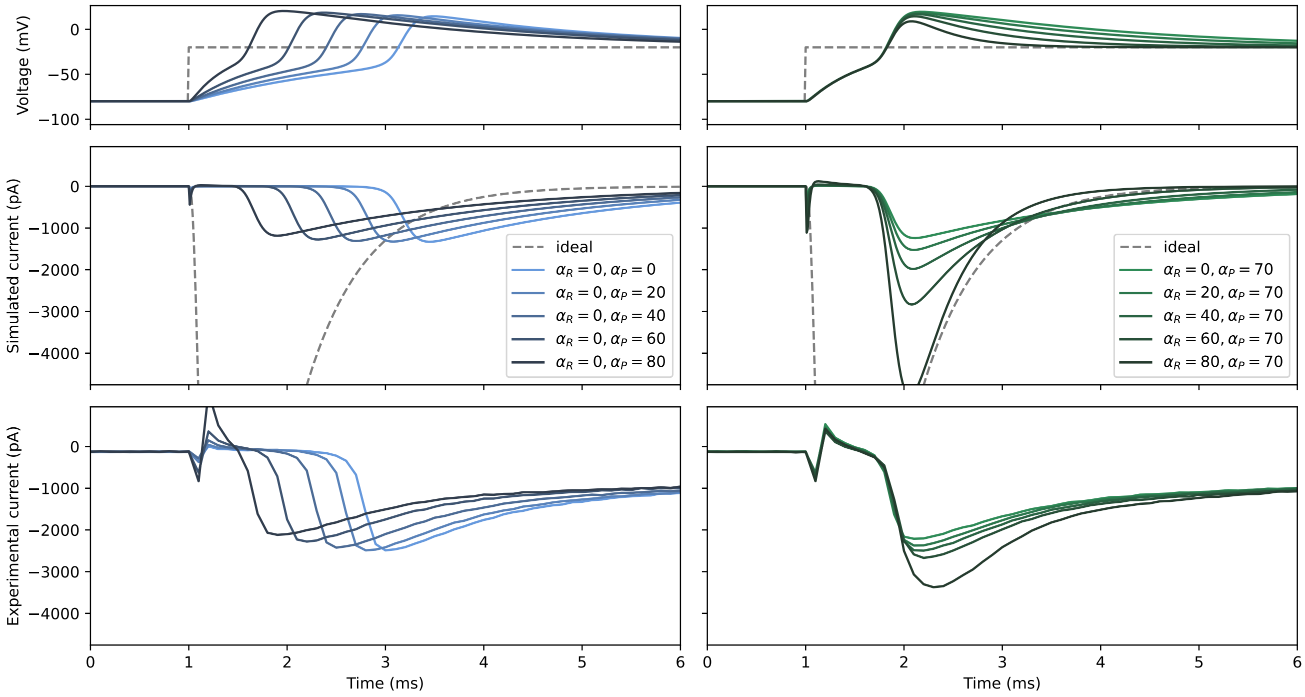


Figure S4: Forward sensitivity analysis of the experimental artefact of fast sodium current using Gray and Franz (2020) and hiPSC-CMs (cell 3).

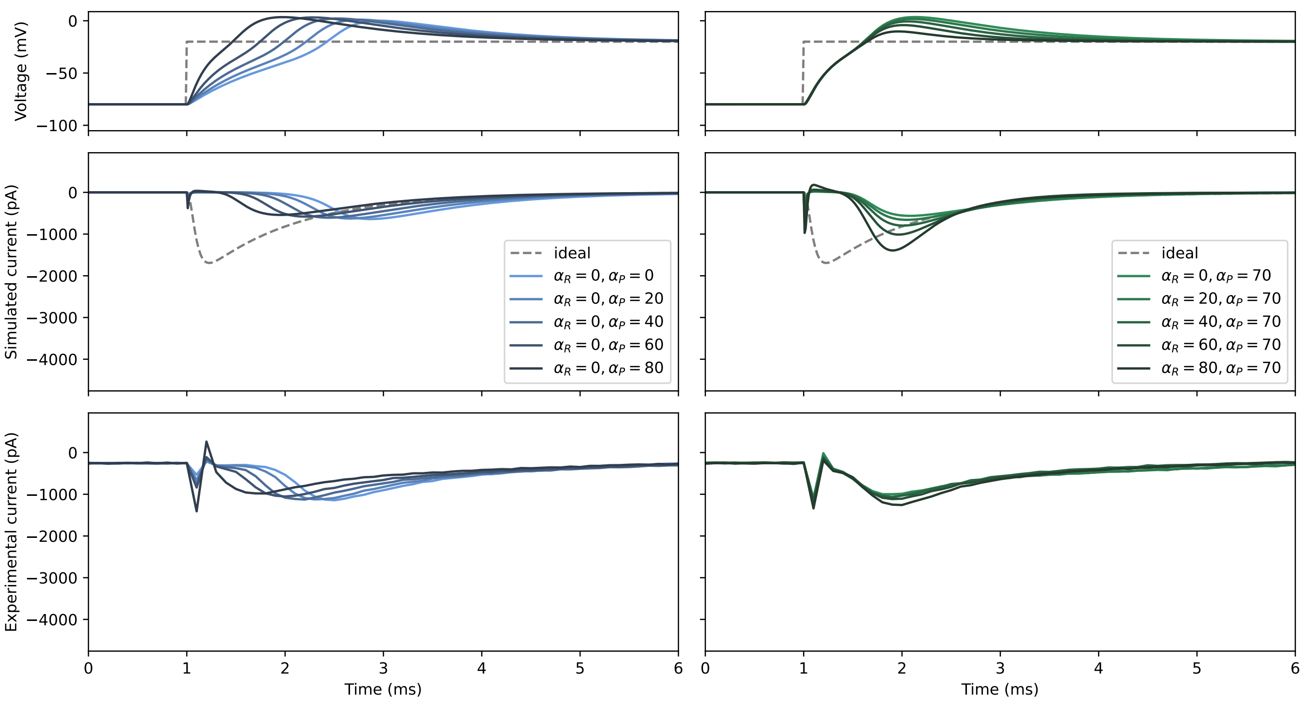


Figure S5: Forward sensitivity analysis of the experimental artefact of fast sodium current using Paci et al. (2020) and hiPSC-CMs (cell 0).

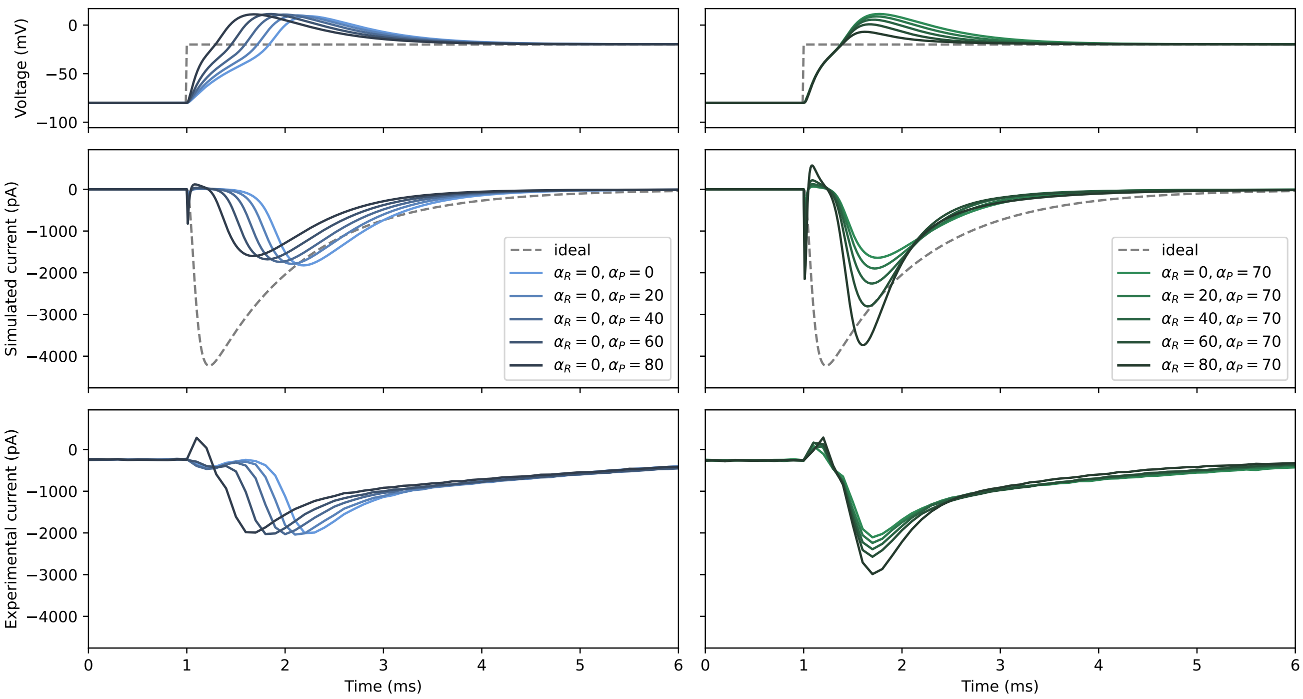


Figure S6: Forward sensitivity analysis of the experimental artefact of fast sodium current using Paci et al. (2020) and hiPSC-CMs (cell 1; same cell as in the main text figure).

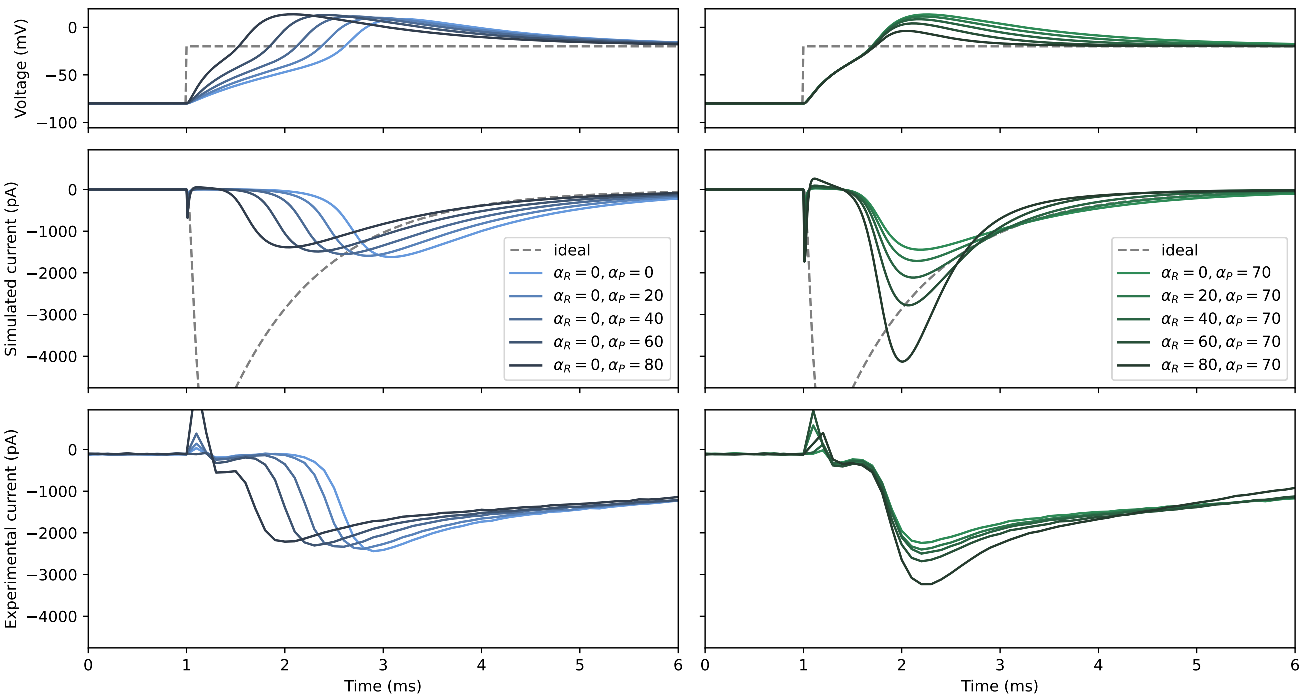


Figure S7: Forward sensitivity analysis of the experimental artefact of fast sodium current using Paci et al. (2020) and hiPSC-CMs (cell 2).

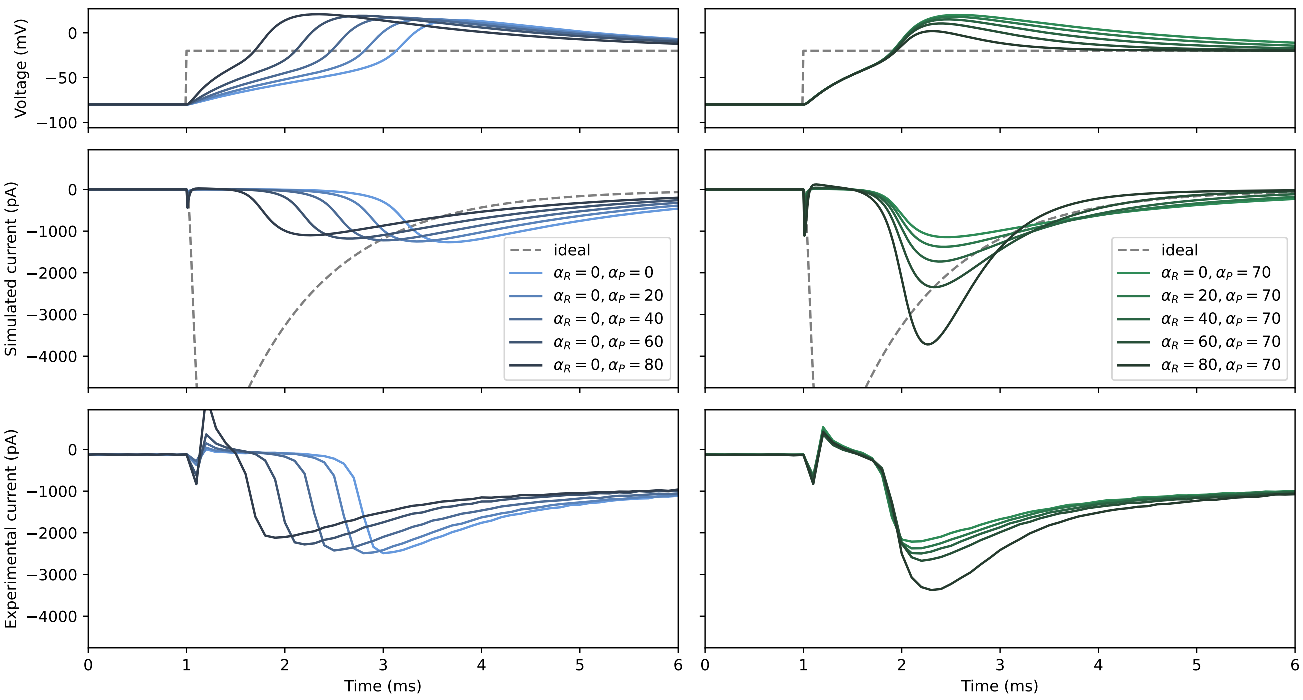


Figure S8: Forward sensitivity analysis of the experimental artefact of fast sodium current using Paci et al. (2020) and hiPSC-CMs (cell 3).

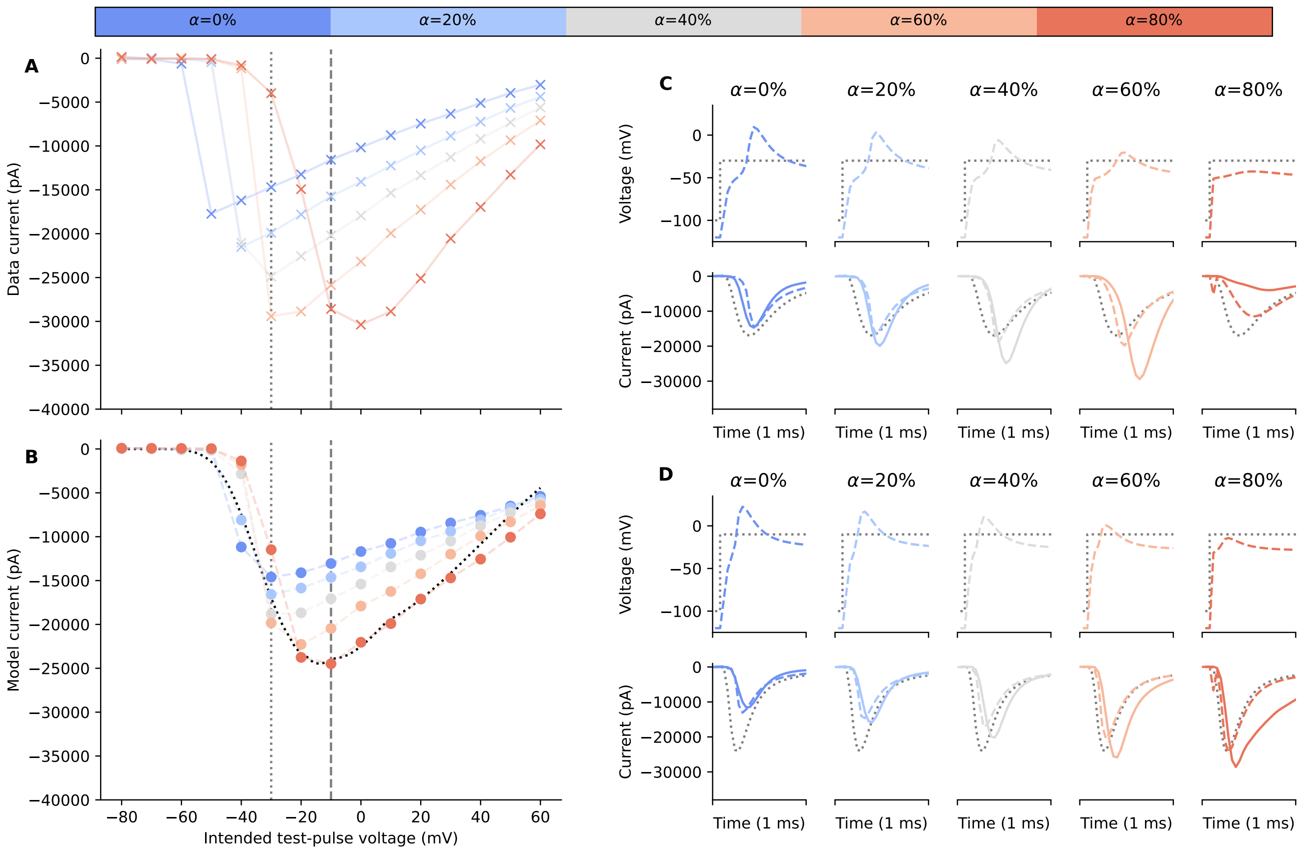


Figure S9: Correction of fast sodium current experiments (cell 1) using the experimental artefact model.

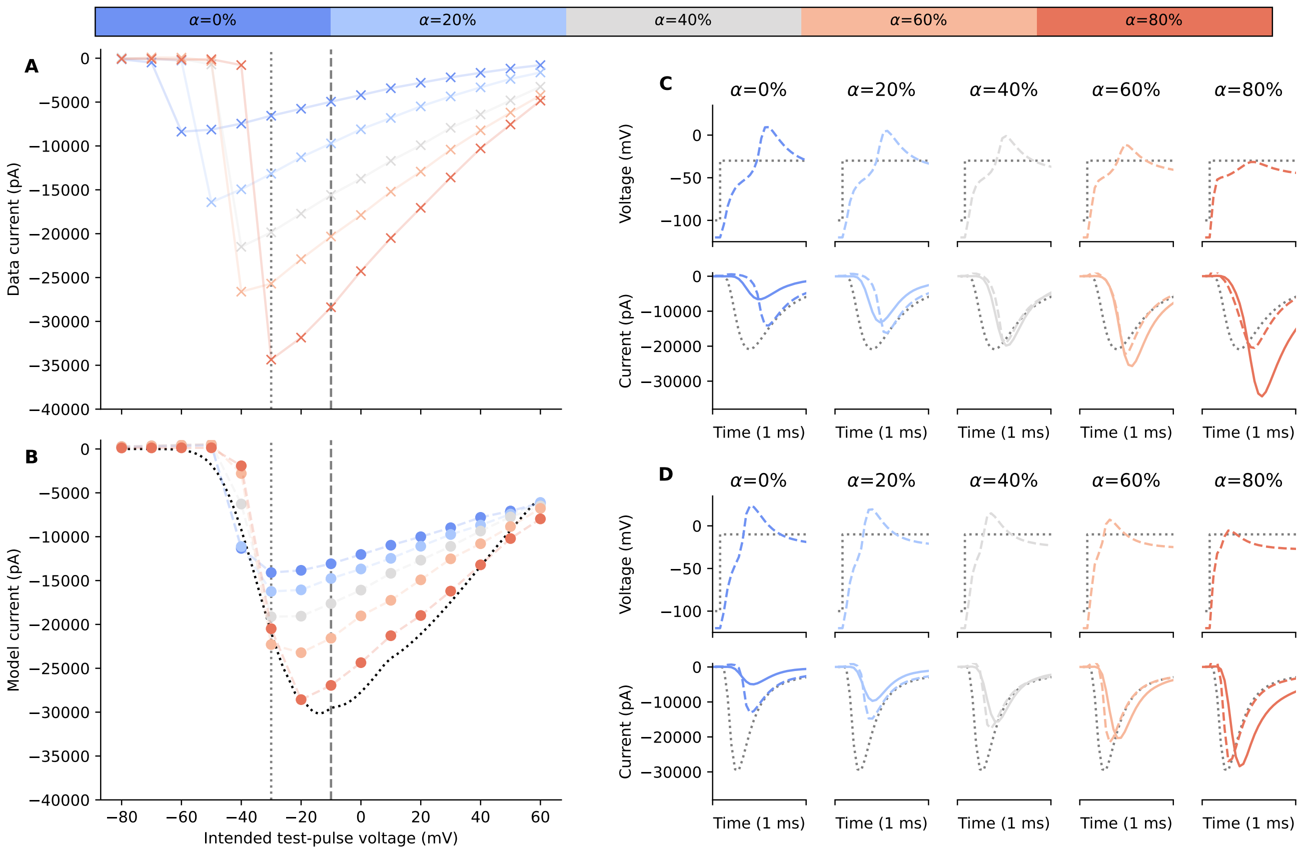


Figure S10: Correction of fast sodium current experiments (cell 2) using the experimental artefact model.

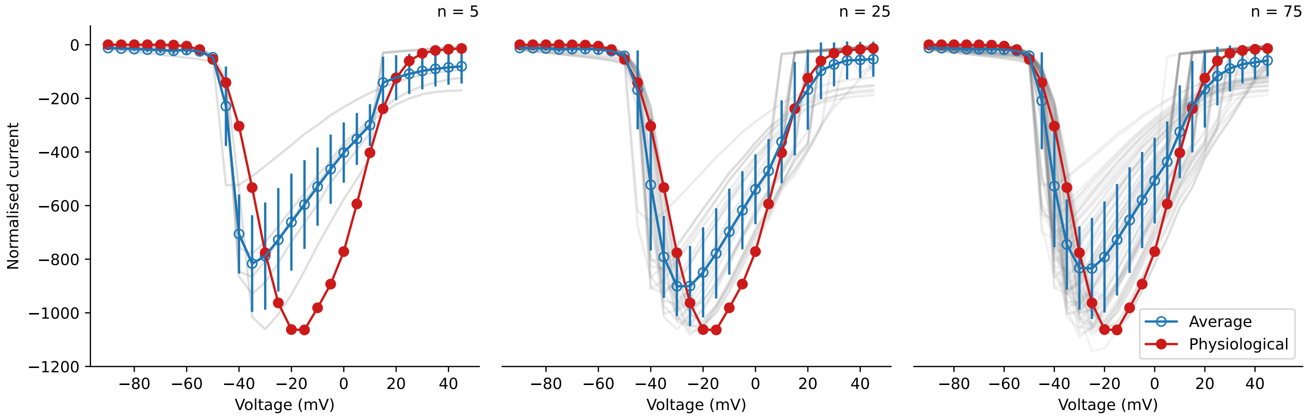


Figure S11: Consequences of averaging fast sodium current normalised I-V curves of multiple runs of experiments, with the average I-V curve shown in blue (averaged from transparent grey lines) and the artefact-free physiological I-V curve shown in red. Left to right shows different number of repeats, $n = 5$, 25, and 75, respectively. Error bars show the standard deviation of the data.

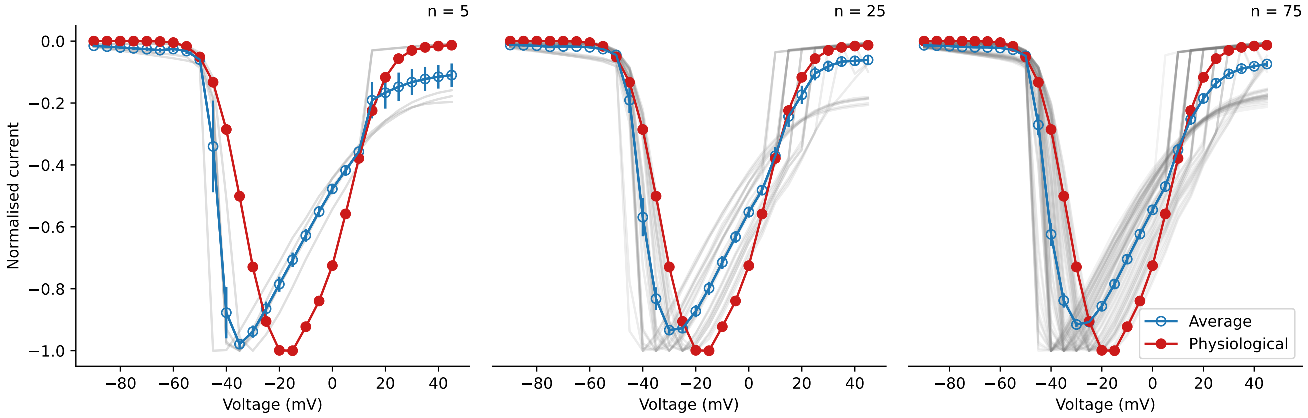


Figure S12: Consequences of averaging fast sodium current normalised I-V curves of multiple runs of experiments, with the average I-V curve shown in blue (averaged from transparent grey lines) and the artefact-free physiological I-V curve shown in red. Left to right shows different number of repeats, $n = 5$, 25, and 75, respectively. Error bars show the standard error of mean (SEM).
